# Differentiation of Pluripotent Synthetic Minimal Cells via Genetic Circuits and Programmable Mating

**DOI:** 10.1101/712968

**Authors:** Nathaniel J. Gaut, Jose Gomez-Garcia, Joseph M. Heili, Brock Cash, Qiyuan Han, Aaron E. Engelhart, Katarzyna P. Adamala

## Abstract

Synthetic minimal cells, here defined as liposomal bioreactors synthesizing protein, are a recent technology that models the intricate gene and protein networks of live cells, without the noise inherent to natural systems. Here we show a toolset for engineering combinatorial genetic circuits in synthetic cells, where distinct populations of synthetic cells control single-gene components, and those populations are assembled via programmable fusion to construct complex biological pathways. We utilize this technology to demonstrate that progenitor populations can mimic differentiation into new lineages in response to small molecule stimuli or as an effect of “mating” with another synthetic cell population. This provides a practical tool for metabolic engineering and natural pathway study, as well as paving the way towards the construction of live synthetic cells with complex gene pathways and the ability to form distinct lineages.

## Introduction

Synthetic minimal cells, liposomal bioreactors capable of producing proteins, are used as models for a variety of biological processes as well as a tools for developing novel functionalities not possible in live cells.^1^ Synthetic minimal cells have been demonstrated to exhibit a number of behaviors previously attributed exclusively to natural living systems, including predation^2^, communication with other cells^3^ and response to their environment^4,5^.

Despite great recent progress, some functionalities remain elusive in synthetic cell systems, including the ability to control expression of more than six genes at a time^6^ and the ability to differentiate populations of synthetic cells into different lineages. To address those needs, we developed a combinatorial genetic circuit assembly technology, where populations of liposomes have control over single-gene components that can be combined via fusion to construct complex biological pathways.

It has been previously shown that SNARE protein mimics (soluble N-ethylmaleimide-sensitive factor attachment protein /coiled-coil hybrid receptors) can be used to fuse phospholipid vesicles and protein expressing synthetic cells.^7,8^ The biggest disadvantage of the SNARE mimic system is that SNARE proteins are not robustly orthogonal – it is difficult to build a system with more than two pairs of SNAREs in the same experiment. This led to the development of a DNA fusion system, which mimicks the SNARE proteins’ membrane fusion mechanism with duplex DNA. The concept of sequential fusion of compartmentalized bioreactors has also been explored using various other systems, including polymersomes^6^, coacervates and, most extensively, liposomes^9,10^. Here we present implementation of a programmable, fully orthogonal fusion system, used to create complex multi-component genetic circuits in synthetic cells. We demonstrate application of this fusion technology to create lineages of synthetic cells; differentiating “ancestral” populations into independent lineages. The ancestral populations are engineered to be “pluripotent,” differentiating upon sensing of a small molecule signal or upon “mating” with another population of synthetic cells.

The applications of the pluripotent synthetic cell platform are not limited to model systems. Previously, it has been shown that up to five genes can be expressed in bulk cell-free protein expression reactions.^11^ We expand this system to a more biologically relevant size of over ten genes in a pathway, inside a synthetic cell liposome, with the ability to independently control the genes.

Cell-free systems are used as a high-throughput screening platform for the discovery of metabolic engineering pathways.^12^ In order to allow investigation of biotechnologically relevant genetic circuits in synthetic minimal cells, we demonstrate engineering combinatorial pathways while maintaining separate reaction environments until the controlled fusion completes the pathway.

## Results and Discussion

Liposome populations can be fused using SNARE protein mimics^9^ or DNA tags^13,14^, as well as various small molecule fusogenic agents^15^. In liposome-based synthetic cells, genetic circuits can be constructed and activated by fusion of two populations using either fusogenic amphiphiles^16^ or SNARE protein mimics^3^. For our goal of building combinatorial libraries of synthetic minimal cells and constructing genetic pathways with multiple nodes, a highly controllable and programmable fusion system was required. This directed us towards the use of nucleic acid membrane tags, rather than small molecules or lipid additives that act non-discriminatively on all membranes present in the sample. DNA tags provide the highest degree of orthogonality, with over hundreds of possible combinations.

### Fusion of liposomes can be detected by membrane and lumen mixing

Classical methods of investigating liposome fusion – FRET membrane dye assays and lumen mixing assays based on either dye or oligonucleotide mixing – are limited to binary systems (looking at fusion of two populations of liposomes). In order to test the multi-component fusion system, we designed a lumen mixing assay with orthogonal pairs of reporter DNA inside liposomes. The fastest and most cost-efficient method of detecting hybridization between multiple pairs of DNA is a small molecule double-stranded DNA specific dye. Unfortunately, many of these dyes respond nonspecifically to a hydrophobic environment. In the presence of lipid membranes, those dyes are sequestered in the lipid bilayer, producing false positive signal. Among many dyes we have successfully used for detection of duplex formation in solution reactions (**Figure S1**), few retain reliable performance in presence of liposomes (**Figure S2**). Of all dyes tested, we chose Diamond, a proprietary nucleic acid dye. This dye displays the largest fluorescence change in the presence of double-stranded DNA versus single-stranded DNA, and it retains this discrimination in the presence of phospholipid membranes. We demonstrated that multiple events of double-stranded DNA formation can be detected in the presence of liposomes (**Figure S3**). To adapt the previously described DNA-mediated liposome fusion system to synthetic minimal cells, we first established a liposome mixing assay: populations of liposomes encapsulating complementary DNA strands, four pairs total, were prepared with DNA fusion tags anchored to the membrane via cholesteryl tags. Each pair of oligonucleotides was designated with lowercase and capital letters – *Aa, Bb, Cc* and *Dd* (**Table S2**). One population of liposomes was decorated with four oligos *A, B, C* and *D*; this population encapsulated four orthogonal sequences of reporter DNA oligos A’, B’, C’ and D’. Four different populations were then decorated with fusion tag oligos consisting of the other half of the reporter oligo duplex (fusion tag *a* on the surface and reporter a’ inside, fusion tag *b* and reporter b’, fusion tag *c* and reporter c’, and fusion tag *d* and reporter d’). This open-ended system allows testing multiple combinations in parallel, enabling future construction of high-throughput platforms for prototyping gene pathways. Using this lumen mixing essay, we tested four pairs of DNA fusion tags, which allowed five independent synthetic cell populations to be mated in programmable and controllable way. Upon sequential mixing of the liposomes, the membrane mixing was detected using pairs of FRET membrane dyes (**Figure S4**) and lumen mixing was detected with Diamond dye (**Figure S5**). The fusion, both measured as membrane and lumen mixing, occurs if, and only if, two membranes contain a complementary pair of fusion tags (*A* with *a, B* with *b*, etc.). After establishing that the fusion system is truly orthogonal and strictly depends on the presence of DNA fusion tags, we proceeded to demonstrate the utility of this system in constructing combinatorial genetic circuits in liposome-based synthetic minimal cells.

### Genetic pathways can be activated by combinatorial mating of synthetic minimal cell populations

We constructed a simple genetic circuit, with two populations of synthetic cells, both containing a bacterial cell-free protein expression system. One population of synthetic cells contains plasmids with GFP under the control of a T7 promoter and the other population contains T7 RNA polymerase under the control of the endogenous bacterial promoter-σ70 (**Figure 1a-c**). Expression of GFP is only possible upon fusion of both populations. In the simplest example, the “ancestral,” or starting, population carries tags *ABCD*, and can “mate” with populations carrying tags *a, b, c* and *d* – each capital and small letter corresponds to a complementary DNA tag sequence (with fusion pairs *A-a, B-b, C-c* and *D-d*). We detected no cross-talk between tags; only membranes labeled with the complementary fusion tags can successfully fuse and produce GFP signal, and fusion mediated by one pair of DNA tags occurs independently of fusion mediated by another pair (**Fig. 1d**).

**Figure 1.**
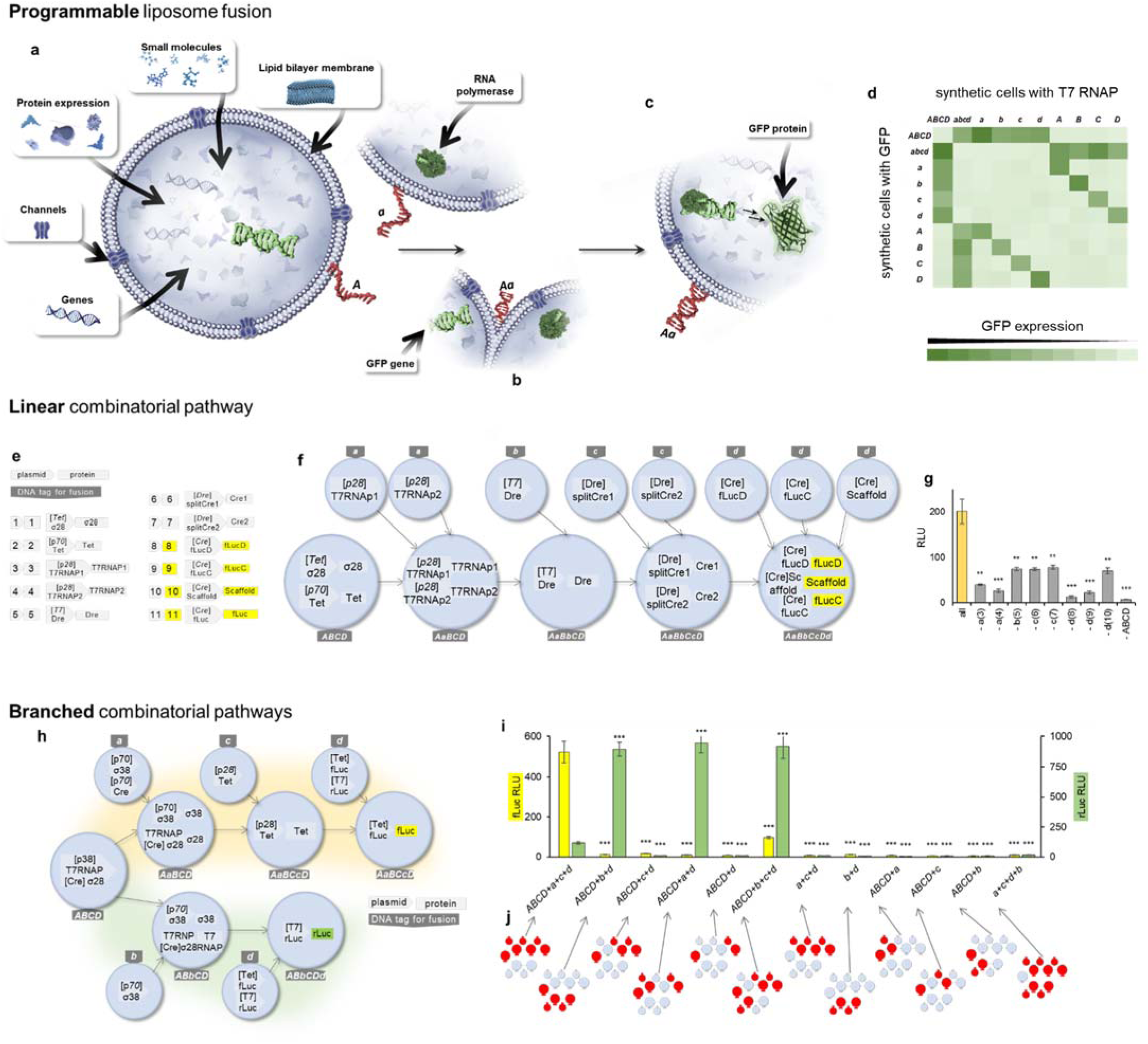
Combinatorial synthetic cell fusion. **a.** Semi-synthetic minimal cell system used in this work. Liposome bilayer encapsulates *E. coli* cell-free protein expression system. **b**. Synthetic cell liposome membranes were decorated with DNA tags (capital and lowercase letters signify complementary pairs). DNA was anchored in the membrane using cholesterol tag. The mechanism of DNA-mediated liposome fusion is analogous to SNARE protein mediated fusion, bringing membranes together by complementary DNA pairs. **c**. Liposome fusion and subsequent combining of the content of two populations of synthetic cells results in activation of genetic circuits. In this example, RNA polymerase from population *a* initiates transcription of gene carried by population *A*, resulting in GFP expression. **d: Mixing of synthetic cells can be detected by construction of a genetic circuit**. Two populations of synthetic minimal cells are decorated with DNA membrane tags. Only if a complementary pair of DNA fusion tags (*A*-*a, B*-*b*, etc.) is present on opposite populations, the membranes can mix, consequently mixing the lumen of the cell as well. One population contains T7 RNA polymerase, the other population contains plasmid for the reporter protein, GFP. GFP expression is measured using fluorescent readout (see **Fig. S10** for individual data points). The heat map represents GFP fluorescence from max 359 to min 14 FU measured at λ_ex_ 488nm and λ_ex_ 509nm. Membrane mixing can also be traced by fusion assays (see **Fig. S4**). **Fusion of synthetic minimal cells containing genetic circuits creates linear (e-g) and non-linear (h-j) combinatorial pathways. e-g**: **A Linear pathway** is constructed with series of RNA polymerases and recombinases. **e**: symbols used on panel **f**, with corresponding shorthand numbers used in panel **g**. Arrows indicate promoters, squares indicate protein products. **f**: genes encapsulated in different populations of synthetic minimal cells. Sequential fusion of populations creates the final circuit; readout of the pathway is the activity of firefly luciferase reconstituted from the three-partite split reporter. The detailed schematic of the pathway is on **Figure S6. g**: Linear genetic pathway. Synthetic cells with membranes decorated with DNA tags were mixed with ancestral synthetic cell population *ABCD*. The content of the fusing liposomes, not the specific identity of the membrane tags, influences the final population state (**Fig. S11**). Variants of those pathways were first tested via simultaneous expression in one solution, without encapsulation in synthetic cells (**Fig. S7**). Variant pathways of varying complexity were created based on this pathway (**Fig. S26**). **h**-**j: Non-linear genetic pathway**. The outcome of the ancestral population *ABCD* depends on the identity and the order of the fusions of the populations. **h**: the schematics of the non-linear pathway with two possible outcomes. **i**: the firefly (yellow) and *Renilla* (green) luciferase signals measured in the final progeny synthetic cell. Error bars signify S.E.M. with n=4. **j**: shows the same synthetic cell populations as panel **h**, with the specific populations used in each experiment highlighted in red.

### Complex linear and branched pathways can be constructed by mating populations of synthetic cells

Synthetic minimal cells have been used as a tool for studying expression of natural gene pathways as well as a model system for creation of synthetic pathways.^17–19^ We asked if the combinatorial mating of synthetic cell populations could produce novel pathways of a size and complexity approximating that of natural biological pathways. Synthetic cells have been successfully used to construct and investigate pathways of up to six genes (albeit using polystyrene compartments instead of phospholipid bilayer membranes).^6^ The most commonly investigated gene circuits in synthetic cells are usually not longer than two to four genes, while biologically relevant gene pathways are typically at least two to three times that size. Additionally, biological pathways are often non-linear, with the output of the pathway depending on activity of genes interacting with each other in sub-networks or branches.

Having established that the DNA tag mediated fusion system allows combinatorial programmable mating of synthetic minimal cells with phospholipid membranes, we proceeded to applying this technology to construct combinatorial genetic pathways of a complexity approaching natural systems. First, we designed and validated a ten gene pathway, constructed from a series of RNA polymerases, recombinases, and a multicomponent luciferase reporter (**Fig. S6**). All variants of this pathway produce a luciferase signal only if all genes are expressed, and modifications of that pathway allow synthetic genetic pathways to be built with varying length and complexity (**Fig. S7**). The linear pathway (**Fig. 1e**) can be constructed by using the DNA mediated combinatorial fusion of synthetic cells: with ancestral population *ABCD* (the populations are named after the fusion tags on the surface of synthetic cells) being mated with eight different populations of synthetic cells decorated with complementary fusion tags (**Fig. 1f**). Only if all populations are mixed and contain the appropriate fusion tags, is the final product of the pathway, luciferase signal, produced (**Fig. 1g**).

To demonstrate a more biologically interesting example of a non-linear pathway, we modified the model luciferase pathway to produce two possible outputs: firefly or *Renilla* luciferase (**Fig. 1h**). This branched pathway started with an ancestral synthetic cell population *ABCD*, and the first fusion event sets the “fate” of the pathway: fusing with population *a* or with population *b*, sets the synthetic cell lineage on a course to produce either firefly or *Renilla* luciferase (**Fig. 1i**). We tested multiple variants of mating scenarios, confirming that only the presence of all necessary fusion partners results in detection of the final pathway product (**Fig. 1j**). We use the term “lineage” to describe a synthetic minimal cell population capable of producing certain output: differentiation from an ancestral population.

### Genetic pathways can be independently activated by mating different components to the same ancestral population of synthetic cells

We found that the identity of populations contributing to the development of any new lineage, and not the order in which the differentiation events occurred, defined the identity of the final lineage. For example, the ancestral population containing T7 and σ28 promoters, can be differentiated into progeny lineages expressing comparable levels of the three measured fluorescent proteins, regardless of the order in which the populations contributing to that lineage were mated (**Figure 2**, the three compared lineages are highlighted by bold rectangles on panels **d**-**f**, a schematic of the circuit on panel **a** and data on panel **b**). Signal measured at the end of each experiment corresponds to the expected level of fluorescence (comparing experimental data on **Figure 2d-f** with theoretically predicted levels of fluorescence on **Figure S8**), regardless of the order in which the elements of the genetic circuit were delivered.

**Figure 2.**
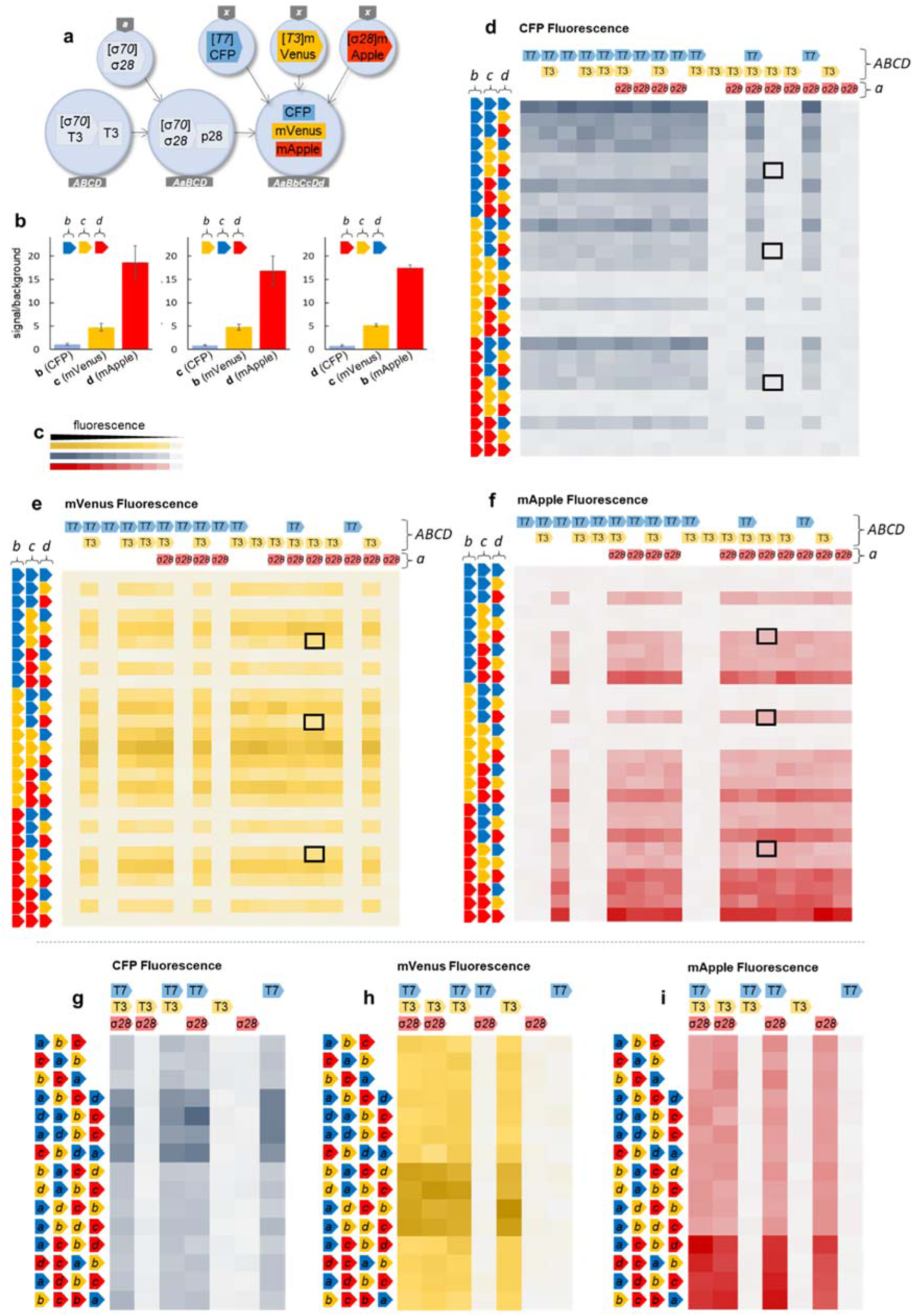
Independent genetic pathways can be activated by combinatorial fusion of different components to the same ancestral population of synthetic cells. The ancestral population *ABCD* contains one or two RNA polymerases (T7 and/or T3). The first fusion event, with synthetic cell population *a*, adds another RNA polymerase to the circuit (σ28), or in case of an empty space, Population *a* contains no plasmid. The order of fusion is always the same, first Population *a*, then *b*, then *c* and then *d*. In each experiment, Populations *b, c* and *d* carry different reporter protein, marked by the color of arrows to the left of each heat map (CFP as blue, mVenus as yellow and mApple as red). This creates a sequence of fusion events, with each reporter protein introduced at a different time. Colors of labels for T7, T3 and σ28 correspond to the colors of reporter proteins controlled by promotors for each polymerase (T7 in blue for T7 promoted CFP, T3 in yellow for T3-promoted mVenus and σ28 in red for σ28-promoted mApple). **a**: A specific example of one of the pathways, the sample indicated by thick black square on panels **d-f**; the ancestral synthetic cell population contained T7 RNAP, the first fusion event brought σ28 polymerase under the endogenous σ70 promotor, and subsequent fusion populations brought CFP under T7, mVenus under T3 and mApple under σ28. **b**: Fluorescent readout from pathway described on panel **a**, presented as ratio of measured fluorescent signal to background in that channel; error bars are S.E.M. with n=3. **c**: Relative intensity scale bar for all data on panels **d-i**. Absolute fluorescence values are in figures **S12, S13, S14, S15, S16** and **S12**. **d-f**: The RNA polymerases in the ancestral populations are labeled on top of the heat map. Each heat map shows fluorescence in separate channel, corresponding to one of the three reporter proteins; **d**: CFP (blue) with λ_ex_ 433nm and λ_em_ 475nm, **e**: mVenus (yellow) with λ_ex_ 515nm and λ_em_ 530nm, **f**: mApple (red) with λ_ex_ 568nm and λ_em_ 592nm. Each data point is the average of 3 experiments. Data for each specific experiment are on Fig. **S12** for the blue, **S13** for yellow and **S14** for the red reporter protein. The observed data fits with the simulated expression profiles, calculated based on the amount and ratio of reporter protein vectors and presence of specific promotors (Fig. **S8**). **g-i**: **The order of fusion events of each subsequent population into the ancestral population does not affect the final outcome**, measured by three different fluorescent protein reporters in the final population. In each experiment, the ancestral population *ABCD* and subsequent populations *a, b, c* and *d* were fused in the order represented by the order of the arrows on the left of each heat map. The ancestral population *ABCD* contains from one to three RNA polymerases (labeled on top of each heat map as T7, T3 and σ28), and subsequent populations *a, b, c* and *d* contain plasmids for reporter proteins CFP (blue), mVenus (yellow) or mApple (red). Data for each specific experiment are on Fig. **S15** for the blue, **S16** for yellow and **S17** for the red reporter protein. The outcomes of the pathways can be compared to the theoretically predicted activity values shown on figure **S9**. All reactions were incubated for 8h post fusion, then liposomes were lysed, protein expression was stopped with the inhibitor cocktail (see Materials and methods for details), and fluorescence of the reporter proteins was measured (CFP at λ_ex_ 433 nm and λ_em_ 475nm, mVenus at λ_ex_ 515nm and λ_em_ 530nm, mApple at λ_ex_ 568nm and λ_em_ 592nm).

### Strength of pathway activation depends on the copy number of each allele

In cell-free translation systems, the strength of gene expression is proportional to the copy number of the gene, within a certain range of DNA concentrations.^20,21^ We observed that this principle is conserved in the genetic circuits assembled in our combinatorial synthetic cell systems. The identity of the final population depends on the copy number of the alleles of each gene, i.e., a progeny population created by the fusion of two populations carrying a gene for a fluorescent protein results in higher signal than fusion of one copy of the given gene if the correct RNA polymerase was present in the ancestral population (**Figure 2g-i**). The observed pathway activation is consistent with the theoretical prediction: we plotted expected fluorescence intensity in each progeny population solely based on the copy number of each gene and the presence of the correct RNA polymerase (**Figure S9**). The results match the patterns observed in the experiments (**Figure 2g-i**).

### Pluripotent synthetic minimal cells give rise to new lineages

To investigate if the differentiation of the ancestral pluripotent synthetic cell population can result in irreversible specific differentiation into separate lineages, we have designed synthetic cells with genomes that can undergo irreversible differentiation events. The differentiation can be either upon detection of a small molecule signal (**Fig. 3a-b**) or upon mating with different populations of synthetic cells carrying distinct differentiation gene signals (**Fig. 3c-i and 3k-r**).

**Figure 3.**
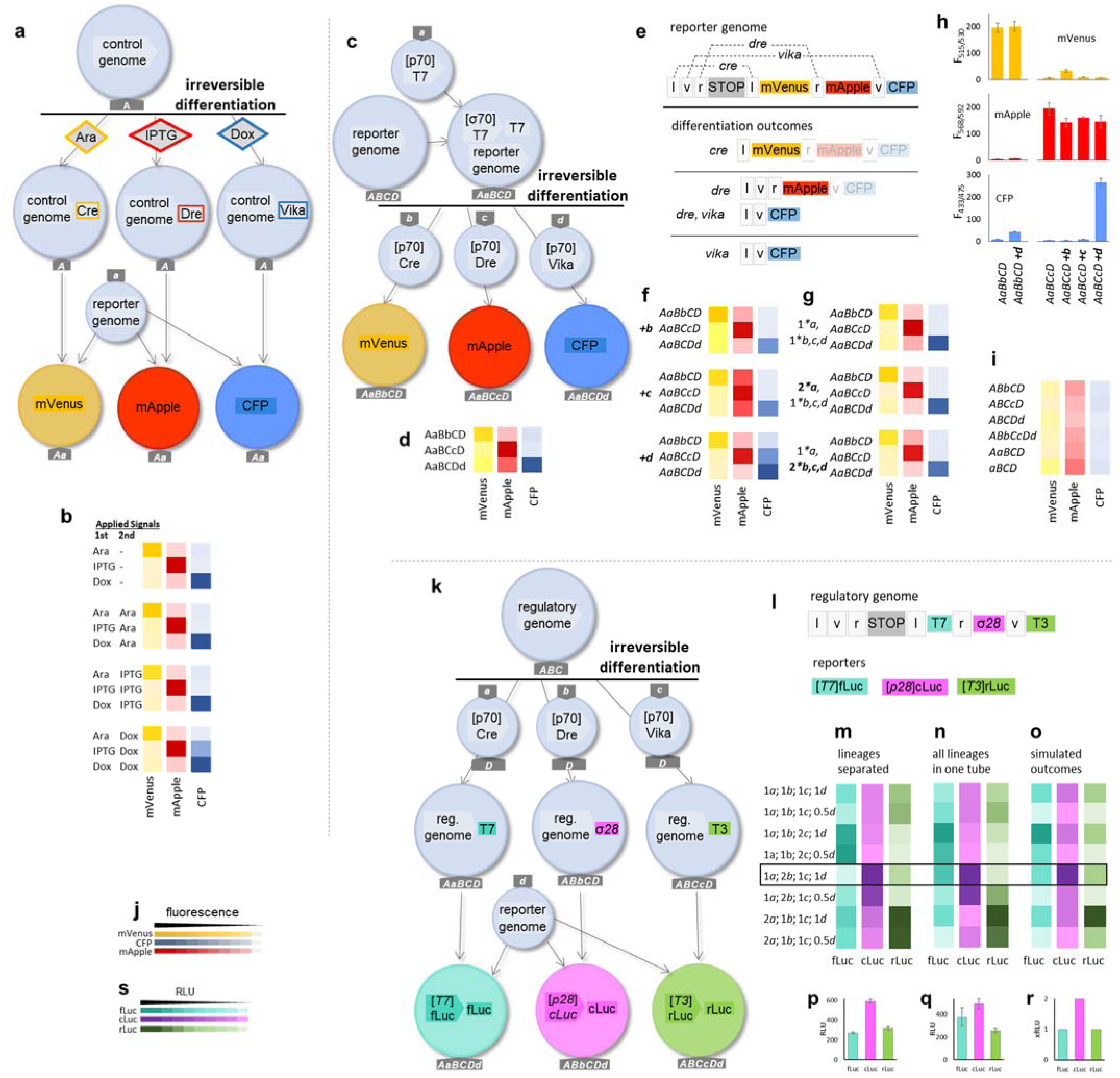
Differentiation of pluripotent synthetic cells. Starting population of synthetic cells can be differentiated into separate lineages in specific and irreversible differentiation events that cause the rearrangements of the genome of the ancestral synthetic cell population. **a-b: Differentiation in response to small molecule signals; a**: This synthetic cell population is pluripotent – it can differentiate into three different lineages, depending on exposure to small molecules. The control genome contains arabinose inducible Cre recombinase, IPTG inducible Dre recombinase and tetracycline (Dox) inducible Vika recombinase. The differentiation step is the induction of expression of the recombinase. Each of the lineages is then fused with a population of synthetic cells containing the reporter genome (shown on panel **e**). **b**: Fluorescence of all samples at the completion of each experiment was measured in three fluorescent channels (depicted as yellow for mVenus, red for mApple and blue for CFP). The resulting lineages express fluorescent reporter protein based on the identity of recombinase acting on the reporter genome (shown in panel **e**). Additional incubation with another signaling molecule does not produce more or different reporter signal for Dre and Cre populations. Additional expression of Vika in originally Dre population produces an increase in CFP signal. Data for each specific experiment are on **fig. S18**. **c-i: Differentiation induced by fusion of different populations of cells. c**: The pluripotent population *AaBCD*, arising from the first fusion event of ancestral population *ABCD* with population *a*, is then irreversibly differentiated by fusion with either population *b* (carrying Cre, resulting in *AaBbCD*, yellow), population *c* (carrying Dre, resulting in *AaBCcD*, red) or population *d* (carrying Vika, resulting in *AaBCDd* blue). **d**: Reporter activity was measured from each new lineage after differentiation, in yellow (mVenus), red (mApple) and blue (CFP) channels. **e:** The reporter genome for experiments outlined in panel **a** and panel **c** contains three fluorescent proteins, flanked by recombinase sites and each open reading frame ends with stop codon. Activity of the recombinase defines the fluorescent protein expressed in any given lineage. Recombination sites: v – Vox site for Vika, r – Rox site for Dre, l – lox site for Cre. **f-i**: **Further differentiation events do not produce additional lineage differentiation. f**: Adding one more equivalent of differentiating populations (+*b*, +*c* or +*d*) to the lineages obtained after initial speciation event (*AaBbCD, AaBCcD* or *AaBCDd*) does not change the observed reporter signal (compared to the original lineages on panel **d**); except in the case of addition of population *d* (carrying Vika) to red population *AaBCcD*, which results in additional CFP expression. **g:** Addition of multiple equivalents of initial population *a* (1\**a* or 2\**a*), or multiple equivalents of differentiating populations *b, c* or *d* (1\**b,c,d* or 2\**b,c,d*) at the time of differentiation, does not change the reporter protein signal from each lineage, as compared to the original lineages on panel **d. h**: Example data from few lineages: population *AaBbCD* (lineage with Cre), the same population subjected to second differentiation event with additional equivalent of population d (carrying Vika), population *AaBCcD* (lineage with Dre) and the same population with additional one equivalent of differentiating populations *b, c* or *d*. **i**: Without the initial fusion with population *a*, as well as without any recombinase from either of the differentiating populations *b, c* or *d*, no activity is detected from any of the reporter genes. Data for each specific experiment for panels d, f, g and i are on **fig. S19**. **j**: Fluorescence heat map scale for panels **b, d, f, g** and **i. s**: Heat map for luciferases activity in panels **m, n** and **o**. Heat maps are relative; in each panel the values are scaled independently, from maximal to minimal value in each sample set. Specific numerical values are available on corresponding SI figures. **k-s: Differentiation of ancestral synthetic cell population can be controlled entirely by the identity of cells fused to the ancestral population. k:** The schematic of the differentiation experiments. The starting population contains only the regulatory genome (shown on panel **l**), and differentiation steps involve fusion with populations carrying one of three recombinases and additional fusion tag *D*. The reporter genome can only fuse with populations that underwent one of the differentiation fusion events (acquiring tag *D*). **l:** The regulatory genome contains the same recombination sites as in experiments on panels **a**-**i**, but placed under endogenous bacterial p70 promotor, with luciferase reporter proteins: firefly luciferase (fLuc), *Renilla* luciferase (rLuc) and *Cypridina* luciferase (cLuc). **m**-**o**: Each experiment was performed by mixing different ratios of four different synthetic cell populations, *a* (carrying Cre), *b* (carrying Dre), *c* (carrying Vika) and *d* (carrying reporter genome) to one equivalent of starting population *ABC*. **m**: When experiments for each lineage were carried out in separate tubes. **n**: When all three lineages were incubated together in one tube. **o**: Simulated outcomes, given in expected ratio of luciferase activity, based on ratios of reporter genome and differentiation event populations. **p**-**r**: Example data points for each experiment in the thick black frame on panels **m**-**o**, error bars are S.E.M. n=3. Data for each specific end point experiment are on **fig. S20** for panel **m** and **S21** for panel **n**.

### Synthetic cells differentiate in response to signaling molecules

Small molecule-mediated cellular signaling is crucial for most processes that require coordination between different cells, including differentiation. It has been demonstrated that synthetic minimal cells can send and receive chemical signals using quorum sensing secondary metabolites in liposomes^4,5,8^ and in droplets^22^, as well as protein^23^ and DNA^24^ mediated communication. We asked if our synthetic cell differentiation could be used to engineer pluripotent synthetic cells that can undergo differentiation in response to small molecule signaling. We designed a control genome (**Fig. 3a**), which contained three recombinase genes under the control of small molecule inducible promotors previously demonstrated to be effective in cell-free protein expression system^11^. We also built a reporter genome (**Fig. 3e**), which contained three reporter fluorescent proteins with expression regulated by the three recombinase sites. The ancestral, pluripotent synthetic cell population *A* (decorated with only one type of DNA fusion tag, the A tag) was treated with one of the three signaling molecules, inducing expression of one of the three DNA recombinases (**Fig. 3a**). This was the irreversible differentiating step: the expression of one of the three possible recombinases defined the kind of reporter genome rearrangement the lineage could undergo. Another population of synthetic cells, containing the reporter genome and decorated with DNA fusion tag a, was then mated with the control genome populations *A*. The reporter genome synthetic cells (tagged *a*) mated with all three lineages, and the final product, fluorescent protein signal, depended solely on the type of recombinase present in the control genome population (tagged *A*) differentiated with the signaling molecule. The recombinases induced irreversible rearrangement of the reporter genome, and each lineage produced fluorescent protein reporter signal that corresponded to the expected recombinase rearrangement product: the lineage in which the control genome was treated with arabinose expressed yellow fluorescent protein mVenus, the lineage treated with IPTG produced red mApple and the lineage treated with doxycycline produced blue CFP (**Fig. 3b**). We have demonstrated that the lineage differentiation was irreversible, as additional treatment with a different signaling molecule after the reporter genome fusion and rearrangement did not produce change in the lineage identity. The only exception was doxycycline treatment of the lineage created with IPTG, which started expressing detectable amounts of blue reporter protein CFP in addition to the expected red mApple, suggesting that Vika recombinase rearranged the reporter genome further, and the Dre-mediated recombination was not complete (**Fig. 3b**).

This experiment demonstrates that pluripotent synthetic minimal cells can undergo irreversible, specific differentiation into lineages producing different protein products upon treatment with small signaling molecules, and the identity of the lineages can be controlled and programmed by mating with other populations of synthetic minimal cells.

### The history of the synthetic cell lineages defines the identity of the final synthetic cell population

To demonstrate differentiation upon mating with another population, we designed a reporter genome containing three recombinase sites and three reporter proteins (**Fig. 3e**). This genome was encapsulated inside population *ABCD*. The initial fusion with population *a* is not a speciation event, but it is crucial in maturing each synthetic cell population in preparation for differentiation. In the presence of T7 RNA polymerase, encapsulated by population *a*, the genome of the ancestral population *ABCD* is transcribed, but not translated, due to the stop codon in front of the reporter genes (**Figure 3c**). The three mating partners (tagged *b, c* or *d*) that can induce differentiation into specific lineages contain one of the three recombinases necessary to induce expression from the reporter genome. Upon mating with either of those differentiation populations, the reporter genome (in *AaBCD*) undergoes irreversible rearrangement, setting the synthetic cell population on *a* path to become one specific lineage. The lineage identity can be characterized based on fluorescent protein signal (**Fig 3d**).

### The recombinase induced differentiation step is irreversible

Subsequent mating events with additional differentiating populations, after the initial irreversible differentiation mating, do not change the final lineage identity (**Fig 3f**). This holds true even after multiple mating events (**Fig 3g**): see representative data points for one specific lineage maturation (**Fig 3h**). Absence of any crucial maturation or differentiation events results in lack of any protein signal characteristic of a differentiated lineage (**Fig 3i**). The exception to this rule was that in the presence of additional population *d* (carrying Vika), there was increased blue (CFP) expression due to further reporter genome rearrangement.

We asked if the ancestral population can be truly “naïve” – not capable of differentiation into any lineage until the first differentiation event. To investigate this, we engineered another recombinase based regulatory genome and reporter genome based on three luciferase plasmids (**Fig 3l**). In this case, the ancestral, pluripotent, synthetic cells *ABC* lack the ability to mate with the reporter genome *d*. Only by mating with populations *a, b* or *c*, can specific lineages be created and the incoming differentiating mating partners deliver the fusion tag *D*, necessary for the final mating event with the population carrying the reporter genome (**Fig 3k**). As in previous experiments, further differentiation events do not change the initial lineage. Interestingly, we demonstrated that all lineages can be incubated together, in the same reaction vessel, creating luciferase readouts that predictably match the expected state of the reporter genomes in each lineage (**Fig 3m-r**). The stoichiometry (ratios of synthetic cells mixed in each population) of differentiation and reporter genome mating events does not affect the identity of the final lineage, but it does affect the amount of the final reporter protein produced. For example, using twice as much of the differentiation population *b* yields an increase in expression of *Cypridina* luciferase (cLuc), as shown on aggregated data panels **3m** (for each lineage incubated separately) and **3n** (for all lineages mixed in a single tube), which is in agreement with the theoretical prediction shown on panel **3o**. Individual data points for the highlighted example are shown on panels **3p-r**.

### Synthetic cell lineages can produce different metabolic pathways

After we demonstrated that combinatorial mating of synthetic cells can create new lineages, we asked if this system could be used practically for engineering gene pathways producing small molecule products. We chose the violacein pathway (**Fig. 4a**), a bacterial pigment producing pathway that is well-studied and characterized under a variety of conditions.^25^ All enzymes of this pathway have been expressed in a cell-free translation system^26^, and the final product and intermediates are natural pigments easily identifiable using HPLC (**Fig. 4c**). The order of the enzymes in the pathway defines the final product (**Fig. 4a**). First, we tested the concept of modular combinatorial pathway design by constructing variants of the Vio pathway. The starting synthetic cell population contained the first enzyme of the pathway (VioA), and that population was labeled with DNA fusion tags *BEDC*. Note that the Vio experiments are the only case where we changed letter designation for DNA fusion tags, instead of *Aa, Bb, Cc* and *Dd* we are using *Bb, Ee, Dd* and *Cc* – to match the letter designations of Vio enzymes. Four different synthetic cell populations were prepared containing the four Vio enzymes necessary to complete the pathway and labeled with DNA fusion tags corresponding to the name of the Vio enzyme, VioB produced by population *b*, etc. Upon fusion of populations *b, e, d* and *c* to the starting population *BEDC*, the complete Vio pathway was constructed and the two main products detected were, as expected, violacein and chromopyrrolic acid (**Fig. 4b**). Omitting one or more of the mating populations resulted in the pathway producing different product distribution based on the enzymes present in the final population (**Fig. 4d-g**). Thus, we demonstrated that we can control the type of small molecule produced inside a synthetic cell population by selectively mating synthetic cells containing different elements of the metabolic engineering pathway. Next, we asked if it is possible to use the metabolic pathway as an identifying feature of a synthetic cell’s lineage by implementing the pluripotent synthetic cell differentiation scheme. We combined the mating induced differentiation concept with the ability to control the identity and future mating potential of each lineage using the incoming mating partners (**Fig. 4h-n**). In this experiment, the starting (ancestral) synthetic cell population *AB* contained the first two pathway enzymes: VioA and VioB. The differentiation event was defined as mating with one of the three populations: all containing VioE, and labeled with either DNA fusion tag *C*, tag *D*, or the pair of tags *C* and *D*. This produced three distinct lineages, all containing the same set of enzymes (VioA, Viob and VioE), but different sets of mating DNA tags (*AaBC, ABbD* or *AaBbCD*). This enabled the second round of mating events to be controlled by the identity of the DNA mating tags on each lineage. Two mating partners were made available to each lineage: a synthetic cell population containing VioC and labeled with DNA mating tag *c* as well as a population containing VioD and labeled with mating tag *d* (**Fig. 4h**). Those new mating partners were only able to mate with lineages displaying the appropriate, complementary DNA mating tags (*C* for *c* and *D* for *d*). Therefore, there were three possible final lineages: *AaBCc* containing all enzymes except VioD (**Fig. 4i** and **l**), *ABbDd* containing all enzymes except VioC (**Fig. 4k** and **n**), and *AaBbCcDd* containing all five Vio pathway enzymes (**Fig. 4j** and **m**). The detected distribution of products in each lineage was consistent with the expected product distribution from the enzymes present in each lineage (**Fig. 4i-k**).

**Figure 4.**
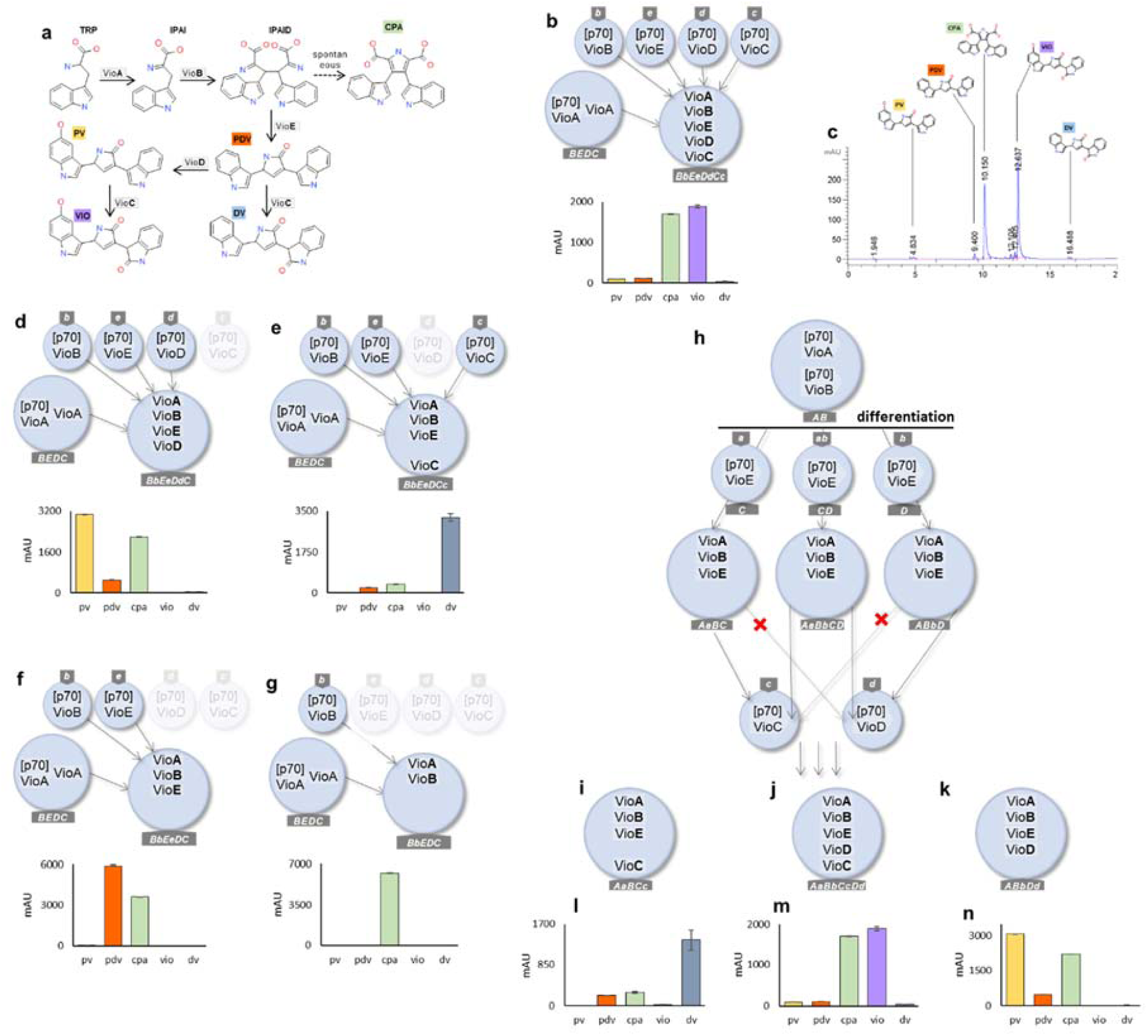
Combinatorial mating of populations of synthetic cells results in construction of divergent variants of an enzymatic pathway. The violaceic acid pathway produces final compounds that depend on the identity and order of the enzymes applied to the pathway intermediates. The synthetic cell mating tags direct the identity of enzymes, and thus the final distribution of the products, in each lineage. On this figure, the annotation of fusion tags has changed: instead of *abcd* -*ABCD* pairs used on previous figures, here the fusion tag letter corresponds to the name of the Vio enzyme carried in each population. The ancestral population always contains VioA, and the fusion populations are labeled *b, e, d* and *c* – corresponding to the Vio enzyme they contain. **a**: the violacein acid pathway; TRP, L-tryptophan; IPAI, indole-3-pyruvic acid imine; IPAID, IPAI dimer; CPA, chromopyrrolic acid; PDV, prodeoxyviolacein; DV, deoxyviolacein; PV, proviolacein; VIO, violacein. The colors used to mark each pathway product used in panel **a** are maintained through all other panels. **c**: One of the replicates of experiment shown on panel **b** (*BbEeDdCc*) as an example chromatogram analysis of the result of the synthetic cell violacein acid pathway reaction. The positions of each compound were confirmed by authentic sample analysis marked on the chromatogram. **b-g**: Starting from synthetic cell population expressing VioA, variants of violacein pathway are constructed by mating the ancestral population with different populations expressing other Vio enzymes. The chemical pathways built in each lineage from panels **b**-**g** is shown on **Figure S22**. Example of HPLC chromatogram for each experiment is shown on **Figure S23**. **h-n**: Pluripotent synthetic cell population can differentiate into lineages producing different products by mating with populations bringing in different fusion tags. The starting population expresses VioA and VioB, with fusion tags *A* and *B*. The differentiation event is fusion between the ancestral population and mating partners bringing in either *C, D*, or *CD* fusion tags (all mating partners express VioE). The resulting lineages can only subsequently mate with populations displaying the appropriate fusion tag. The allowable mating events produce three distinct lineages (panels **i, j** and **k**), each lineage expresses a different set of Vio enzymes and thus produces a different distribution of products (results shown on panels **l, m** and **n** for product distribution in each lineage). All bar graphs represent total integrated area of each peak, error bars are S.E.M. with n=3.

### Summary

Synthetic minimal cells are rapidly gaining popularity as a crucial means for studying natural systems, and engineering novel biological tools. Expanding the capabilities of synthetic cells enables mimicry of complex life-like cellular properties while maintaining the programmability and relative simplicity of *in vitro* systems.

In this work, we have demonstrated a method for stepwise assembly of biological circuits previously inaccessible within cell-free systems due to their complexity and large size (**Fig 1**). We now may investigate gene pathways with comparable complexity to that of natural genetic systems.

We have also demonstrated a programmable mating system that enables construction of progeny synthetic cells from defined starting populations. Additionally, we have engineered pluripotent synthetic minimal cells, in both lineage agnostic and lineage specific systems. We demonstrated that synthetic cell populations can be designed to have identity that is either independent of the route of construction (lineage agnostic, **Fig 2**) or one that is strictly controlled by differentiation events (lineage specific, **Figs 3** and **4**).

The possible applications of this system include studying pathways involved in natural pluripotent cell differentiation, including effects of small molecule signaling. Another area of possible application is in building tools for metabolic engineering and pathway discovery, potentially creating libraries of gene pathways with different enzyme compositions and ratios. In this context, synthetic cells can be thought of as bioreactors, encapsulating artificial enzymatic pathways.

Alternatively, this technology can be utilized as a tool for construction of complex, live synthetic cells, enabling independent control over different parts of cell metabolism using easily deliverable small molecules, or eventually building complex synthetic cell operons and feedback loops.

This work demonstrates that, while synthetic minimal cells still lack robust reproduction mechanism, it is possible to program differentiation events into populations of synthetic cells. With this work we have advanced the construction of life-like synthetic cells, enabling engineering of multi-gene pathways, and programming development of synthetic cell lineages.

## Acknowledgments

We thank Dr Claudia Schmidt-Dannert for gift of Vio pathway plasmid and for helpful discussion about optimizing the pathway. We thank Dr Vincent Noireaux for gift of T7 RNAP plasmid and for helpful discussion about troubleshooting and optimizing cell-free protein expression system. We thank Dr Edward Boyden for gift of several plasmids used for construction of genes used in this work. We thank Dr Michael Freeman for gift of OphA protein from *Omphalotus olearius* Jack-o’-Lantern mushroom. We thank Dr Richard Murray for helpful discussions about liposome formation systems and cell-free protein expression technologies. We thank Dr Lynn Rothschild for helpful discussions about evolution and differentiation in biological systems.

This work was supported by generous gifts from Jeremy Wertheimer, the Hackett Royalty Fund award, NIH 5R01MH114031-02, NSF CON000000070526, NASA CON00000065217, Semiconductor Research Corporation 2018-SB-2837-C and John Templeton Foundation Exploring the Informational Transitions Bridging Inorganic Chemistry and Minimal Life awards.

## Materials and methods

### Cell-free protein expression reagents preparation

This protocol was adapted from Noireaux^27^ and Jewett^28^. A 50 mL starter culture of the Rosetta 2 strain of *E. coli* (Millipore Sigma 71400-3) was made from a glycerol stock and grown to saturation at 37 °C overnight in 2xYPTG (16 g/L tryptone, 10 g/L yeast extract, 5 g/L NaCl, 20 mM glucose, 40 mM potassium phosphate dibasic, 22 mM potassium phosphate monobasic) with chloramphenicol. A 750 mL 2xYPTG culture (without antibiotic) was inoculated with 10 ml of starter culture. The culture was grown at 37 °C to an OD_600_ of 0.4-0.6, then harvested. The resulting pellet was washed twice with 200 mL of Wash Buffer A (10 mM Tris acetate pH 8.2, 14 mM magnesium acetate, 60 mM potassium acetate, 2 mM DTT) followed by a third wash of 40 ml. The resulting cell pellet can be flash frozen and stored at −80 °C prior to the following steps.

Frozen pellets were suspended in cold Wash Buffer A equivalent to 1.1 times the cell mass. The cells were then lysed by sonication at 4°C with a total output energy of 1.7kJ using the following sonicator settings: 50% amplitude, repeating 10 seconds on followed by 15 seconds off. The lysate was centrifuged at 13,000 × g at 4 °C for 30 min. The lysate was split into 500 µL fractions and incubated at 37 °C with shaking for 1 hour and the tube lids ajar. The lysate is then centrifuged at 13,000 × g at 4 °C for 30 min. The lysate was then split into 50 µl single-use aliquots, flash frozen, and stored at −80 °C until use.

The 10x energy mix is composed of the following: 500 mM HEPES, pH 8; 15 mM ATP; 15 mM GTP; 9 mM CTP; 9 mM UTP; 2 mg/ml *E. coli* tRNA; 0.68 mM Folinic Acid; 3.3 mM NAD; 2.6 mM Coenzyme-A; 15 mM Spermidine; 40 mM Sodium Oxalate; 7.5 mM cAMP; 300 mM 3-PGA.

The 10x energy mix is composed of 17 mM of the following amino acids: alanine, arginine, asparagine, aspartic acid, cysteine, glutamic acid, glutamate, glycine, histidine, isoleucine, leucine, lysine, methionine, phenylalanine, proline, serine For a TxTl (transcription – translation, *in vitro* protein expression) reaction the following components are combined in the indicated concentrations: 12 mM Magnesium glutamate; 140 mM potassium glutamate; 1 mM DTT; 1.5 µM T7 RNA polymerase; 0.4 U/µl Murine RNase Inhibitor; 1x of aforementioned energy mix, 1x of the amino acid mix, and cell free prep. Template concentration varies. Unless otherwise specified, the reaction was incubated at 30 °C for 12 hours followed by 4 °C temperature hold. For liposome experiments, all TxTl reactions contained 200mM sucrose.

### Liposome formation

Liposomes from dioleoylphosphatidylcholine (DOPC), dioleoyl-sn-glycero-3-phosphoethanolamine (DOPE), and cholesterol at a molar ratio of 3:1:1 were prepared according to a previously published method^29^. Briefly: thin film was prepared by mixing all lipids in chloroform (in amber vials) and evaporating the solvent overnight. 500 µL of mineral oil was added to each thin film vial, incubated at 60 °C for 10 min, vortexing for 10 minutes, incubating for 60 °C for 3 hours, then sonicating in bath sonicator at 60 °C for 30 minutes. The mineral oil lipid samples were cooled down to 4 °C. 30 µL of internal liposome solution was added to each mineral oil sample, vortexed for 30 seconds, and equilibrated for 10 minutes at 4 °C.

The mineral oil liposome solution was carefully added on top of 250 µL of centrifuge buffer (100 mM HEPES + 200 mM glucose, pH 8). Samples were centrifuged at 18,000 rcf at 4 °C for 15 minutes. As much of the mineral oil layer as possible was removed, and using fresh pipette tip liposome “pellet” was aspirated form the bottom of the tube. The liposomes were resuspended in 250 µL wash buffer (100 mM HEPES + 250 mM glucose, pH 8), and centrifuged at 12,000 rcf at 4 °C for 5 minutes. Residual mineral oil was removed from the top of the solution, and the liposomes were transferred to a fresh tube.

### Lipid mixing assay

The lipid mixing assay was performed as previously described.^8,13,14^ Briefly, the DNA strands were pre-hybridized by incubating in thermocycler, cooling down from 95 °C to 25 °C over the period of 15 minutes. The pre-hybridized DNA fusion tags in 100 mM HEPES, pH 8.0 were added to a solution of pre-formed liposomes. Liposomes were mixed immediately after addition of DNA, and samples were tumbled for 30 minutes to allow DNA incorporation into liposomes.

### Fluorescent protein measurements in synthetic cells

To ensure that the background fluorescence does not change over the course of a TxTl reaction, we have prepared liposome samples with either no template, or control non-fluorescent protein. The control protein was non-fluorescent OphA protein from *Omphalotus olearius* Jack-o’-Lantern mushroom. Expression of the control protein was confirmed on the Western Blot. See **Figure S24**. To confirm that light scattering from liposome particles does not influence fluorescence signal and the fluorescence signal ratios at different wavelengths, we have confirmed that fluorescence signal from synthetic cell populations expressing different fluorescent proteins does not change upon lysing liposomes (data not shown). We have optimized liposome lysis conditions to make sure the liposomes are always completely lysed, using calcein self-quenching assay. (**Figure S25**)

### Luciferase measurements in differentiated synthetic cells

Liposomes of the non-differentiated ancestral population were labeled with 100 µM of Sulfo-Cyanine5 carboxylic acid. Cy5 fluorescence was recorded at λ_ex_ 646nm and λ_em_ 662nm, and it was used to ensure no loss of ancestral liposome population in every experiment.

The background fluorescence of TxTl reaction was measured in negative control sample included with each series of experiments. The negative control sample contained all components of each experiment except plasmids that produce fluorescent protein signal.

CFP fluorescence was measured λ_ex_ 433nm and λ_em_ 475nm, mVenus fluorescence was measured λ_ex_ 515nm and λ_em_ 530nm, and mApple fluorescence was measured λ_ex_ 568nm and λ_em_ 592nm. All measurements were performed with plate reader PMT setting “high”, and with 12 reads per well. All fluorescent measurements were performed on SpectraMax

### Luciferase assays on differentiated synthetic cells

Liposomes of the non-differentiated ancestral population were labeled with 100 µM of Sulfo-Cyanine5 carboxylic acid. Cy5 fluorescence was recorded from all 3 aliquots before addition of luciferase substrates. The recorded values (data not shown) were used to ensure no loss of ancestral liposome population in every experiment.

### Violacein analysis

HPLC peak assignments were based on previously published data^26,30,31^ and confirmed with the original samples of Violacein from *Janthinobacterium lividum* (Sigma Aldrich) and Deoxyviolacein (Cayman Chemicals).

For HPLC analysis, the violacein pathway reagents were extracted from each sample using ethyl acetate, as demonstrated before.^26^ Briefly, each sample was extracted with 5 parts of ethyl acetate per 1 part sample, vortexing for 5 minutes. The phase separation took approximately 20 to 30 minutes, which is much longer than previously reported (approximately 5 minutes for solution TxTl); we attribute this difference to the presence of lipids in the sample. After complete phase separation, each sample was frozen in an ethanol / dry ice bath, and the ethyl acetate phase was collected.

Samples were analyzed on Agilent ZORBAX Bonus-RP 3.5 µm, 4.6 × 150 mm column, with detection at 570nm, flow rate 1 ml/min, with gradient of acetonitrile 5% in water for 5 minutes, up to 95% at 20 min, down to 5% from 30 min to 35 min. Between each run, the column was washed with 95% acetonitrile in water for 10 min and 5% for 5 min.

## Literature

1. Schwille, P. et al. MaxSynBio – Avenues towards creating cells from the bottom up. Angew. Chemie Int. Ed. (2018). doi:10.1002/anie.201802288

2. Qiao, Y., Li, M., Booth, R. & Mann, S. Predatory behaviour in synthetic protocell communities. Nat. Chem. 9, 1–23 (2016).

3. Adamala, K. P., Martin-Alarcon, D. A., Guthrie-Honea, K. R. & Boyden, E. S. Engineering genetic circuit interactions within and between synthetic minimal cells. Nat. Chem. 1–9 (2016). doi:10.1038/NCHEM.2644

4. Lentini, R. et al. Integrating artificial with natural cells to translate chemical messages that direct E. coli behaviour. Nat. Commun. 5, 4012 (2014).

5. Lentini, R. et al. Two-Way Chemical Communication between Artificial and Natural Cells. ACS Cent. Sci. acscentsci.6b00330 (2017). doi:10.1021/acscentsci.6b00330

6. Peters, R. J. R. W. et al. Cascade reactions in multicompartmentalized polymersomes. Angew. Chemie – Int. Ed. 53, 146–150 (2014).

7. Heo, P. et al. Green fluorescence protein-based content-mixing assay of SNARE-driven membrane fusion. Biochem. Biophys. Res. Commun. 1–7 (2017). doi:10.1016/j.bbrc.2017.05.006

8. Adamala, K. P., Martin-Alarcon, D. A., Guthrie-Honea, K. R. & Boyden, E. S. Engineering genetic circuit interactions within and between synthetic minimal cells. Nat. Chem. 9, 431–439 (2017).

9. Meyenberg, K., Lygina, A. S., van den Bogaart, G., Jahn, R. & Diederichsen, U. SNARE derived peptide mimic inducing membrane fusion. Chem. Commun. 47, 9405 (2011).

10. Ishikawa, K., Sato, K., Shima, Y., Urabe, I. & Yomo, T. Expression of a cascading genetic network within liposomes. FEBS Lett. 576, 387–390 (2004).

11. Shin, J. & Noireaux, V. An E. coli cell-free expression toolbox: Application to synthetic gene circuits and artificial cells. ACS Synth. Biol. 1, 29–41 (2012).

12. Contreras-Llano, L. E. & Tan, C. High-throughput screening of biomolecules using cell-free gene expression systems. Synth. Biol. 3, 1–13 (2018).

13. Stengel, G., Zahn, R. & Höök, F. DNA-induced programmable fusion of phospholipid vesicles. J. Am. Chem. Soc. 129, 9584–9585 (2007).

14. Löffler, P. M. G. et al. A DNA-Programmed Liposome Fusion Cascade. Angew. Chemie – Int. Ed. 13228–13231 (2017). doi:10.1002/anie.201703243

15. Estes, D. J., Lopez, S. R., Fuller, a O. & Mayer, M. Triggering and visualizing the aggregation and fusion of lipid membranes in microfluidic chambers. Biophys. J. 91, 233–43 (2006).

16. Caschera, F. et al. Programmed vesicle fusion triggers gene expression. Langmuir 27, 13082–13090 (2011).

17. Harris, D. C. & Jewett, M. C. Cell-free biology: Exploiting the interface between synthetic biology and synthetic chemistry. Curr. Opin. Biotechnol. 23, 672–678 (2012).

18. Sun, Z. Z., Yeung, E., Hayes, C. A., Noireaux, V. & Murray, R. M. Linear DNA for rapid prototyping of synthetic biological circuits in an Escherichia coli Based TX-TL Cell-Free System. ACS Synth. Biol. (2013).

19. Niederholtmeyer, H. et al. Rapid cell-free forward engineering of novel genetic ring oscillators. Elife 4, 1–18 (2015).

20. Noireaux, V., Bar-Ziv, R. & Libchaber, A. Principles of cell-free genetic circuit assembly. Proc. Natl. Acad. Sci. U. S. A. 100, 12672–12677 (2003).

21. Shin, J. & Noireaux, V. Efficient cell-free expression with the endogenous E. Coli RNA polymerase and sigma factor 70. J. Biol. Eng. 4, 8 (2010).

22. Dupin, A. & Simmel, F. C. Signalling and differentiation in emulsion-based multi-compartmentalized in vitro gene circuits. Nat. Chem. 11, (2019).

23. Niederholtmeyer, H., Chaggan, C. & Devaraj, N. K. Communication and quorum sensing in nonliving mimics of eukaryotic cells. Nat. Commun. 9, 1–8 (2018).

24. Joesaar, A. et al. DNA-based communication in populations of synthetic protocells. Nat. Nanotechnol. 14, 369–378 (2019).

25. Durán, N. et al. Violacein: properties and biological activities. Biotechnol. Appl. Biochem. 48, 127 (2007).

26. Pardee, K. et al. Portable, On-Demand Biomolecular Manufacturing. Cell 167, 248–259.e12 (2016).

27. Sun, Z. Z. et al. Protocols for Implementing an Escherichia coli Based TX-TL Cell-Free Expression System for Synthetic Biology. J. Vis. Exp. 1–15 (2013). doi:10.3791/50762

28. Kwon, Y. C. & Jewett, M. C. High-throughput preparation methods of crude extract for robust cell-free protein synthesis. Sci. Rep. 5, 1–8 (2015).

29. Fujii, S. et al. Liposome display for in vitro selection and evolution of membrane proteins. Nat. Protoc. 9, 1578–1591 (2014).

30. Lee, M. E., Aswani, A., Han, A. S., Tomlin, C. J. & Dueber, J. E. Expression-level optimization of a multi-enzyme pathway in the absence of a high-throughput assay. Nucleic Acids Res. 41, 10668–10678 (2013).

31. Balibar, C. J. & Walsh, C. T. In Vitro Biosynthesis of Violacein from L -Tryptophan by the Enzymes VioA – E from Chromobacterium V iolaceum †. Biochemistry 45, 15444–15457 (2006).

