## Supplementary material for "Differentiation of Pluripotent Synthetic Minimal Cells via Genetic Circuits and Programmable Mating": SI materials

### Figure S1

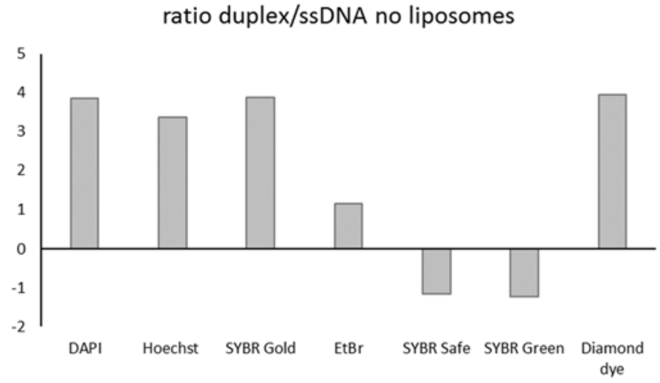


**Figure S1 Testing different dyes for the detection of hybridization of DNA in solution.**
Ratio of dye signal in presence of duplex to single stranded DNA for most popular DNA dyes. Experiment conditions: 25 mM tris-HCl pH 7.5, 100mM KCl, 10 mM MgCl2, 1uM DNA 1 (working stock of 50uM), 5 times addition of 0.2uM DNA2 (working stock of 10uM). All dyes are used at concentration 0.5x, where x is the concentration suggested by manufacturer to use in gel experiment. In many cases, because of proprietary formulation, we are unable to determine the exact molarity of the dye solution.
Reported values are arithmetical average of three replicates.

The fluorescence of the samples was measured at the wavelength specific to each dye:

| dye | $\lambda_{ex}$ (nm) | $\lambda_{em}$ (nm) |
|---|---|---|
| Thiazole orange | 512 | 530 |
| DAPI | 341 | 452 |
| Hoechst | 350 | 461 |
| Ethidium bromide | 510 | 605 |
| SYBR Gold | 495 | 537 |
| SYBR Green II | 497 | 520 |
| SYBR Safe | 502 | 530 |
| Diamond dye | 495 | 558 |

### Figure S2

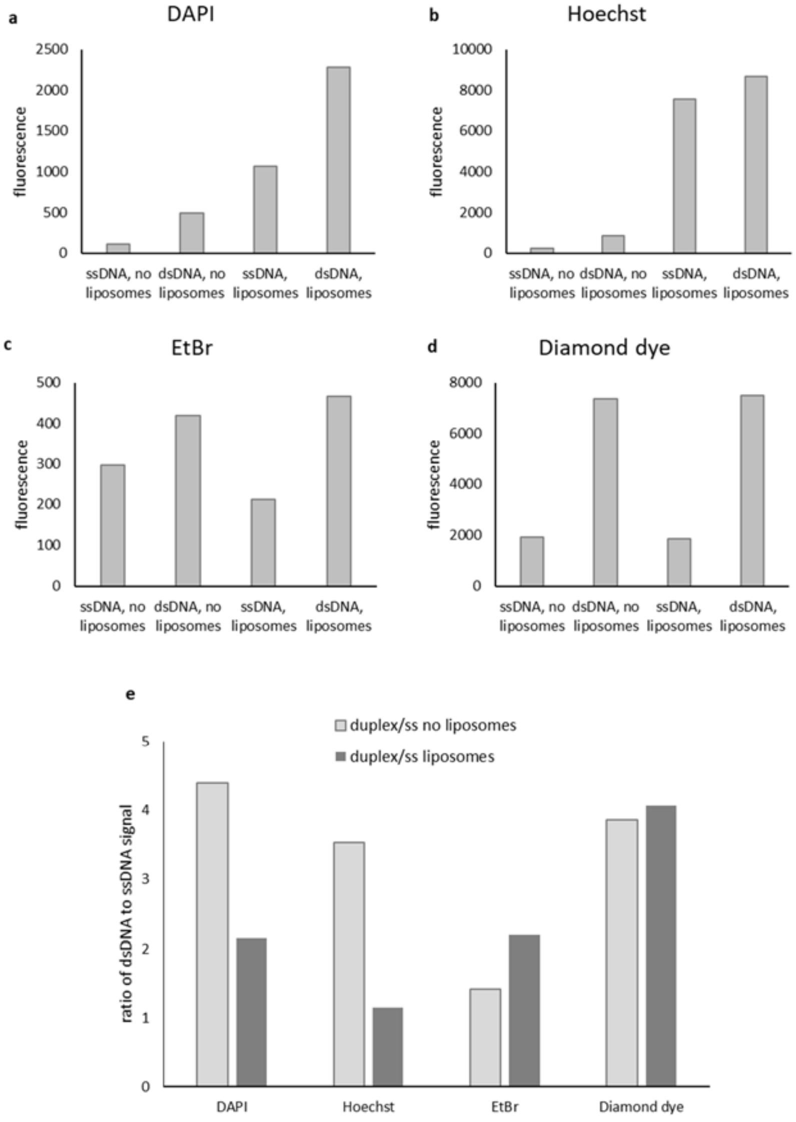


**Figure S2. Testing different dyes for the detection of hybridization of DNA in presence of phospholipid liposomes.**

DAPI (a), Hoechst (b), Ethidium bromide (c) and Diamond dye (d) were used to measure fluorescence of single and double stranded DNA in presence and absence of liposomes. e: comparison of ratio of duplex to ssDNA signal for all four tested dyes, in the presence and absence of liposomes.

Solution experiment conditions: 25 mM tris-HCl pH 7.5, 100mM KCl, 10 mM $MgCl_2$, 1uM DNA 1 (working stock of 50uM), 5 times addition of 0.2uM DNA2 (working stock of 10uM).

Liposome experiment conditions: 25 mM tris-HCl pH 7.5, 5mM, 100mM KCl, 10 mM $MgCl_2$, 1uM DNA 1 (working stock of 50uM), 5 times addition of 0.2uM DNA2 (working stock of 10uM).

All dyes are used at concentration 0.5x, where x is the concentration suggested by manufacturer to use in gel experiment. In many cases, because of proprietary formulation, we are unable to determine the exact molarity of the dye solution.

Reported values are arithmetical average of three replicates.

The fluorescence of the samples was measured at the wavelength specific to each dye:

| dye | excitation wavelength(nm) | emission wavelength(nm) |
|---|---|---|
| DAPI | 341 | 452 |
| Hoechst | 350 | 461 |
| Ethidium bromide | 510 | 605 |
| Diamond dye | 495 | 558 |

### Figure S3

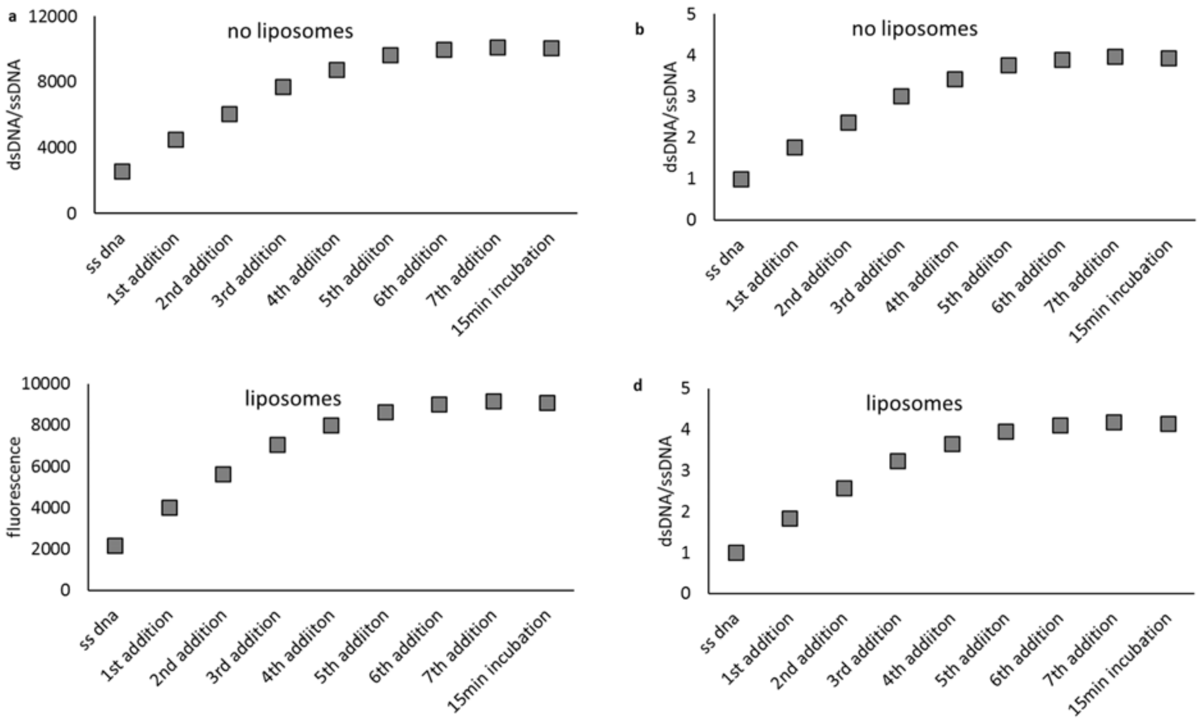


**Figure S3. Sequential duplex formation by addition of DNA to the Diamond dye sample in solution and in presence of liposomes.**

DNA duplex formation measured by fluorescence (a) and represented as ratio of dsDNA to ssDNA (b) in the absence of liposomes, and the same process measured by fluorescence (c) and represented as ratio of dsDNA to ssDNA (d) in the absence of liposomes.

Solution experiment conditions: 25 mM tris-HCl pH 7.5, 100mM KCl, 10 mM MgCl2, 1uM DNA 1 (working stock of 50uM), 5 times addition of 0.2uM DNA2 (working stock of 10uM).

Liposome experiment conditions: 25 mM tris-HCl pH 7.5, 5mM liposomes, 100mM KCl, 10 mM MgCl2, 1uM DNA 1 (working stock of 50uM), 5 times addition of 0.2uM DNA2 (working stock of 10uM).

Reported values are arithmetical average of three replicates.

### Figure S4

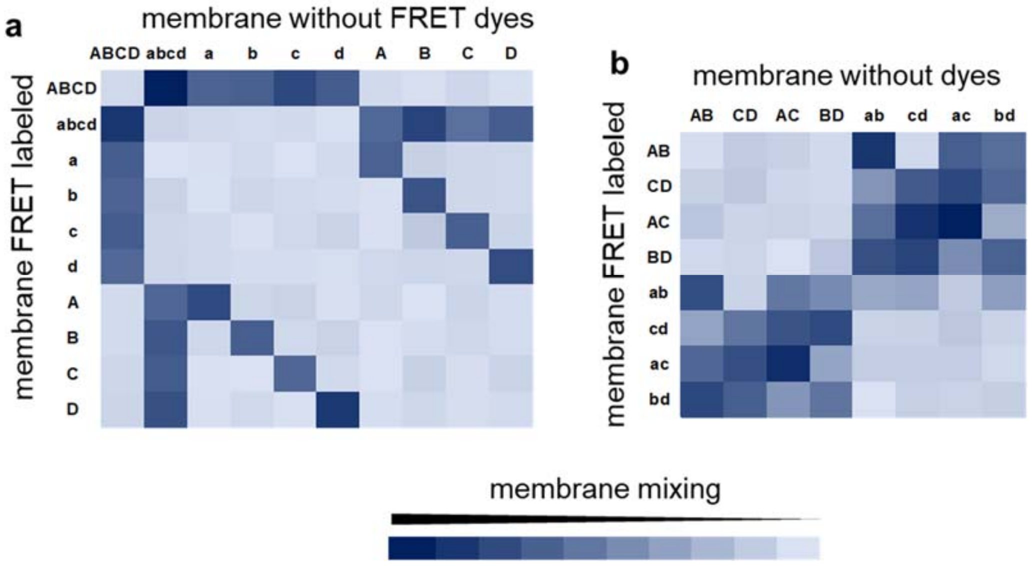


**Figure S4. Mixing of liposome membranes can be tracked by FRET assay, via FRET dyes attached to the membrane of one of the populations.**
Only if complementary pair of DNA fusion tags (A-a, B-b, C-c, or D-d) is present on opposite populations, the membranes mix. Upon mixing, the total surface area of the new liposome is bigger, resulting in increased distance between FRET dyes (see **Figure S27** for schematic of the FRET membrane mixing essay).
**a**: liposome with FRET dye pairs labeled with all four, or every single one of the possible DNA strands, were mixed with liposomes with unlabeled membrane (see **Figure S28** for individual data points).
**b**: same conditions as **a**, but liposomes were labeled with pairs of DNA (see Fig. **S29** for individual data points).
Theoretical calculation was done to predict the outcome of the membrane mixing assays, see **Figure S30**.

### Figure S5

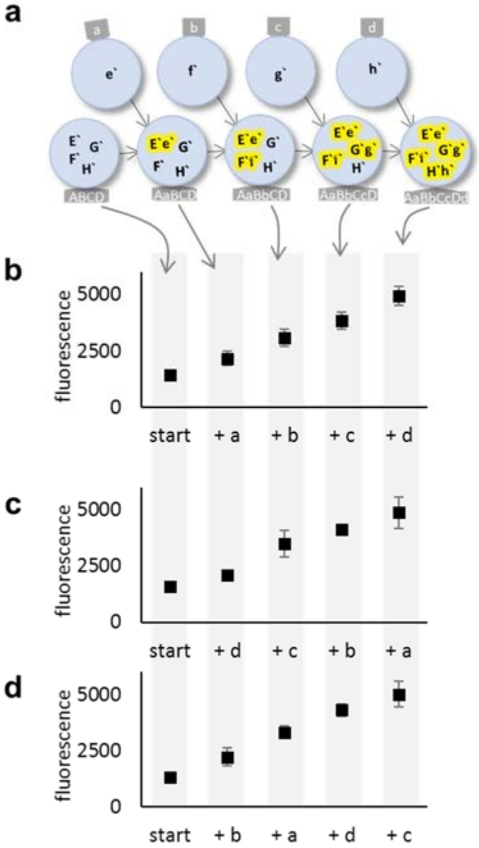


**Figure S5 Liposome content mixing can be detected with a DNA dye that increases fluorescence when ssDNA becomes dsDNA**.

With each mixing event, the dye fluorescence increases. The dye was selected for high ds/ssDNA selectivity (**Fig. S1**) and detection in presence of phospholipid membranes (**Fig S2** and **S3**). **a**: Liposomes are decorated with complementary pairs of DNA anchored into the membrane, A-a, B-b, C-c and D-d. Inside liposomes, complementary DNA pairs E`-e`, F`-f`, G`-g` and H`-h` form duplexes upon mixing of lumen of liposomes. **b - d**. Order of mixing of liposomes does not affect efficiency of content mixing after each event.

### Figure S6

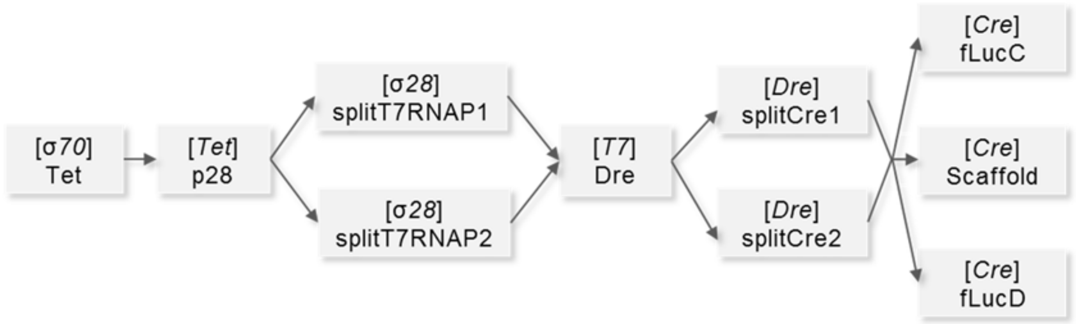


**Figure S6. The schematic representation of the linear multicomponent genetic pathway used as a model pathway in this study.**
Model 10-protein gene pathway designed for optimizing design of multicomponent pathways in synthetic minimal cells. Tet protein is expressed under endogenous P70 promotor, then Tet activates expression of σ28 transcription factor, which turns on two genes under P28 promotor: the halves of split T7 RNA polymerase; upon reconstitution of functional T7 the Dre recombinase is expressed and it induces recombination to excise stop codon and allow expression of two split halves of Cre recombinase; then reconstituted functional Cre turns on expression of three genes: two halves of split luciferase (fLucC and fLucD) that bind to a third protein: a common template (scaffold)[1].

### Figure S7

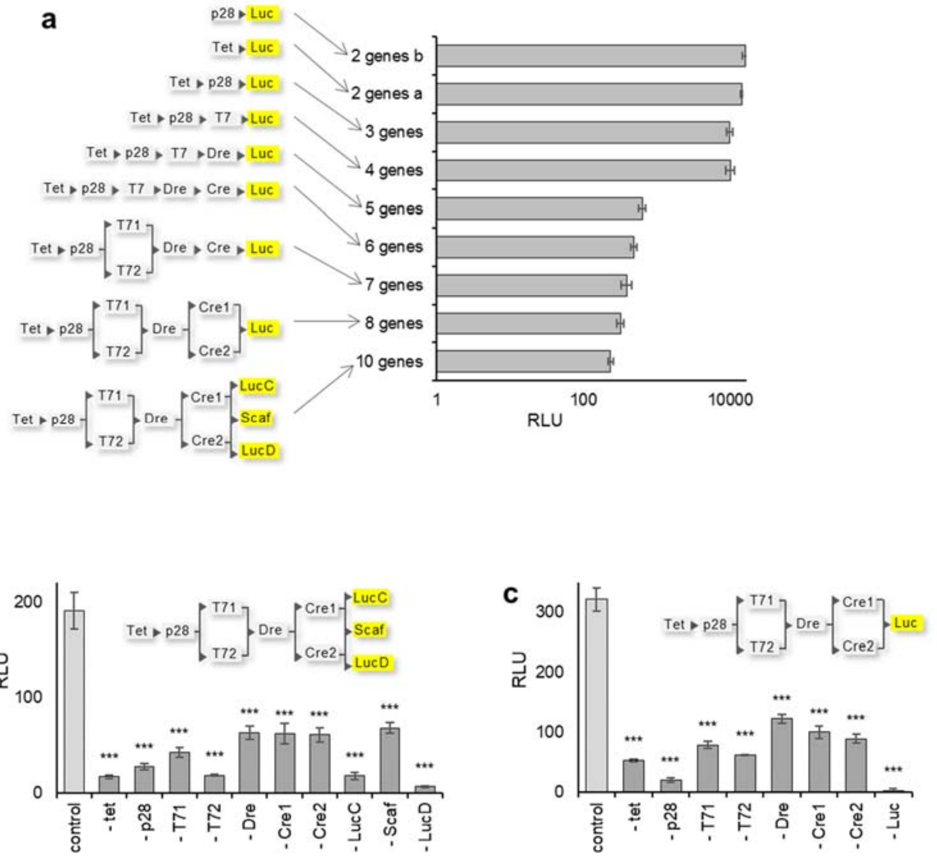


**Figure S7. Testing modularity of the model pathways.**
**a**: Several variants of the model pathway were tested, including complete pathway (from **Fig**. **S6**), and shorter analogues of the full pathway.
**b** and **c**: The modularity is demonstrated on two variants of the model pathway; only if all genes necessary for completing the pathway are present, the final luciferase product can be detected.
All genes were mixed in solution (without liposomes), and they were incubated for 12h. Error bars are ±S.E.M., n=3.

### Figure S8

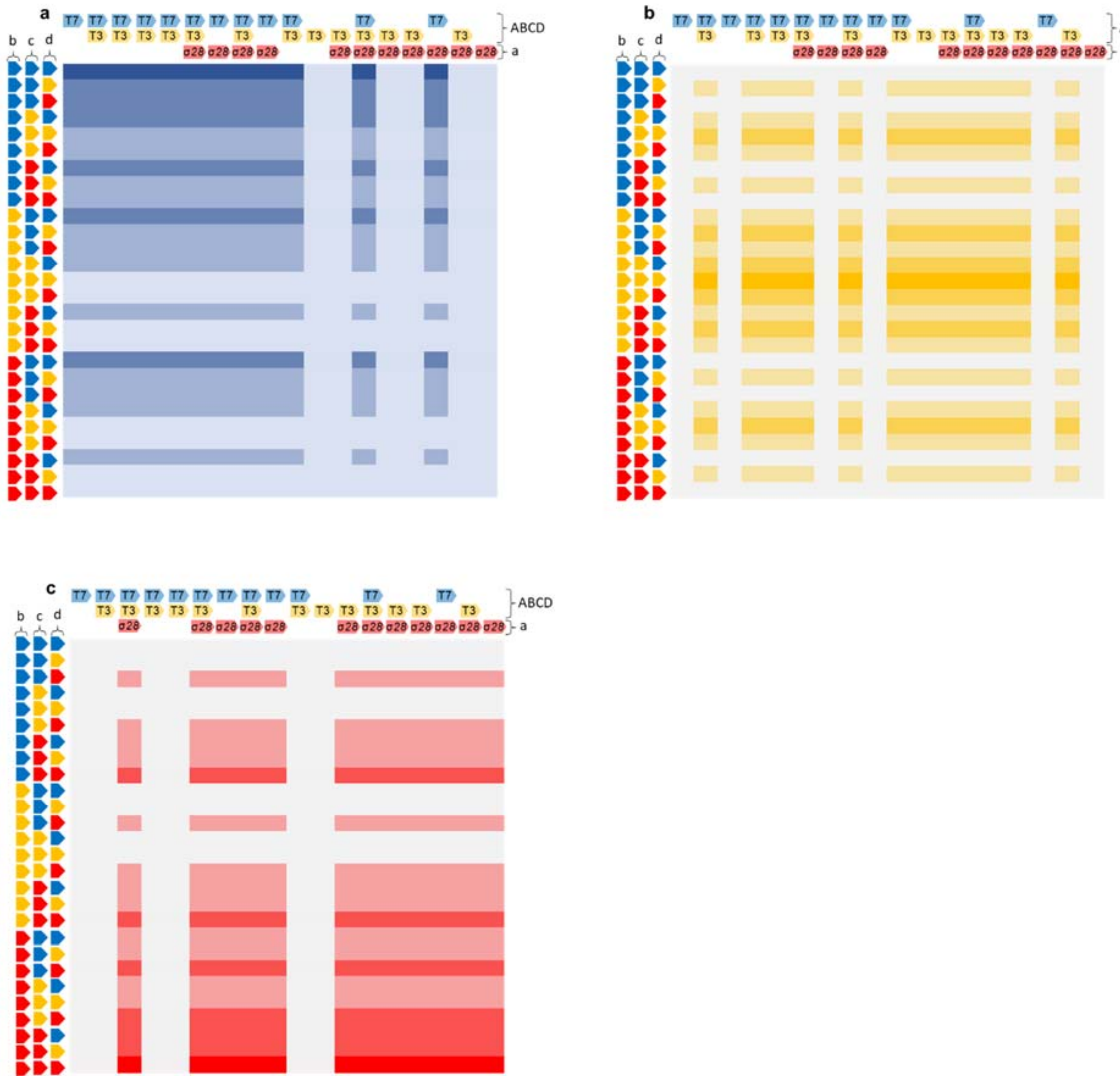


**Figure S8** Simulated activity of combinatorial genetic circuits, calculated based on the amount and ratio of reporter protein vectors and presence of specific promotors.

The ancestral population ABCD contains one or two RNA polymerases (T7 and/or T3), labeled on top of each heat map. First fusion event, with synthetic cell population a, adds another RNA polymerase to the circuit (σ28), or in case of empty space population a contains no plasmid. Three subsequent fusion events, with synthetic cell populations b, c and d, add three reporter proteins to the circuit. The order of fusion is always the same, first population a, then b, then c and then d. In each experiment, populations b, c and d carry different reporter protein, marked by the color of arrows to the left of each heat map (CFP blue, mVenus yellow and mApple red). This creates sequence of fusion events, with each reporter protein introduced at different time. Colors of labels for T7, T3 and σ28 correspond to the colors of reporter proteins controlled by promotors for each polymerase (T7 blue ofr CFP, T3 yellow for mVenus and σ28 red for mApple). The expected outcomes of each pathway was simulated by assigning value of 0 to

fluorescence from sample containing no reporter gene and/or no promotor for that reporter gene. Value of 100 was assigned if sample contained one copy of reporter gene with its cognate promotor, value of 200 was assigned for two copies of that reporter gene and value of 300 for three copies.
Each heat map shows expected fluorescence in separate channel, corresponding to one of the three reporter proteins; **a**: blue CFP, **b**: yellow mVenus, **c**: red mApple.

### Figure S9

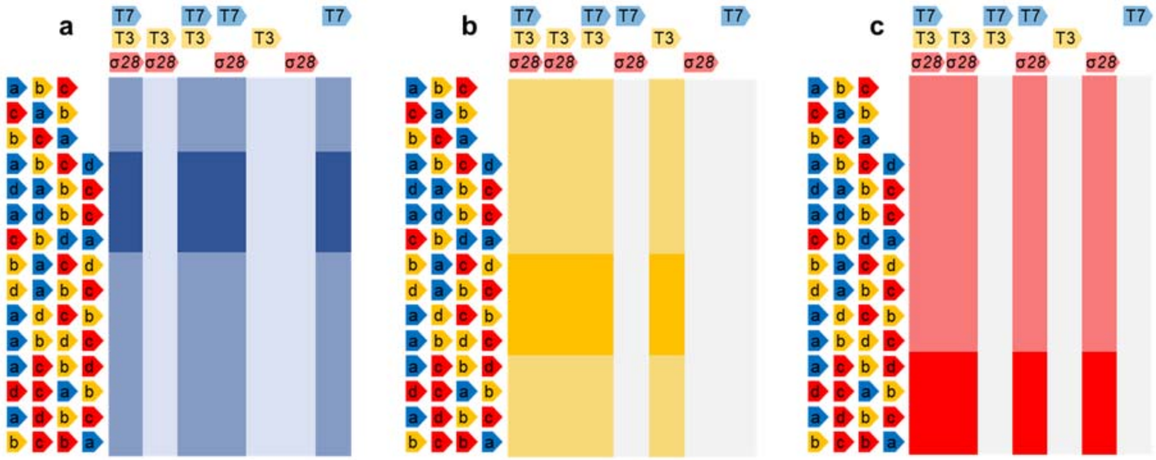

**Figure S9 Simulated activity of combinatorial genetic circuits, calculated based on the amount and ratio of reporter protein vectors and presence of specific promotors.**
In each experiment, the ancestral population ABCD and subsequent populations a, b, c and d were fused in the order represented by the order of the arrows on the left of each heat map. The ancestral population ABCD contains from one to three RNA polymerases (labeled on top of each heat map as T7, T3 and σ28), and subsequent populations a, b, c and d contain plasmids for reporter proteins CFP (blue), mVenus (yellow) or mApple (red). Experimental data from this system are on **Figure 2f-h**.
The expected outcomes of each pathway was simulated by assigning value of 0 to fluorescence from sample containing no reporter gene and/or no promotor for that reporter gene. Value of 100 was assigned if sample contained one copy of reporter gene with its cognate promotor, value of 200 was assigned for two copies of that reporter gene and value of 300 for three copies. Each heat map shows expected fluorescence in separate channel, corresponding to one of the three reporter proteins; **a**: blue CFP, **b**: yellow mVenus, **c**: red mApple.

### Figure S10

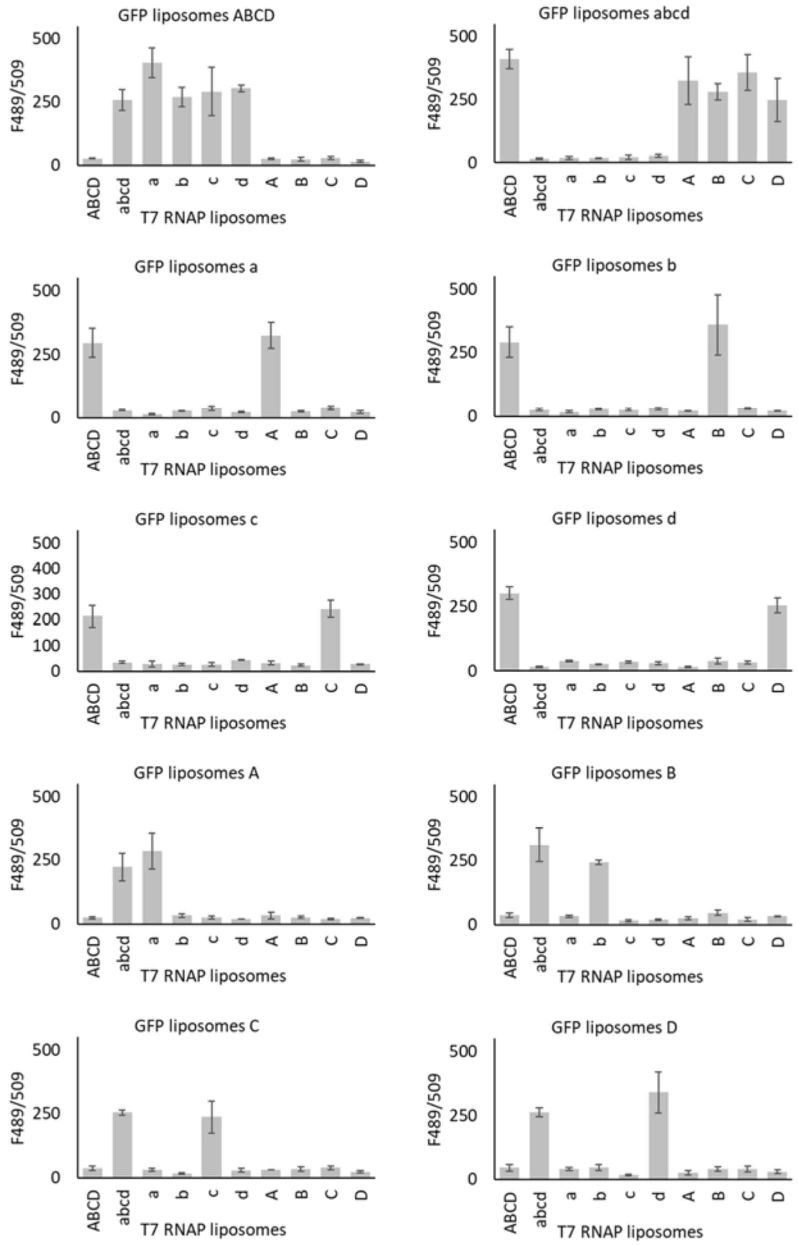


**Figure S10 Individual data points for lumen mixing experiments for synthetic minimal cells labeled with membrane anchored DNA fusion tags.**

Liposomes were prepared with either all (ABCD, abcd) or one of the complementary DNA oligos (a, b, c, d, A, B, C or D). One population of liposomes contained T7 RNA polymerase, the opther population contained eGFP reporter protein. Liposomes were mixed in equimolar amounts, incubated for 12h, and fluorescence was measured. Error bars indicate SD, n=3.

### Figure S11

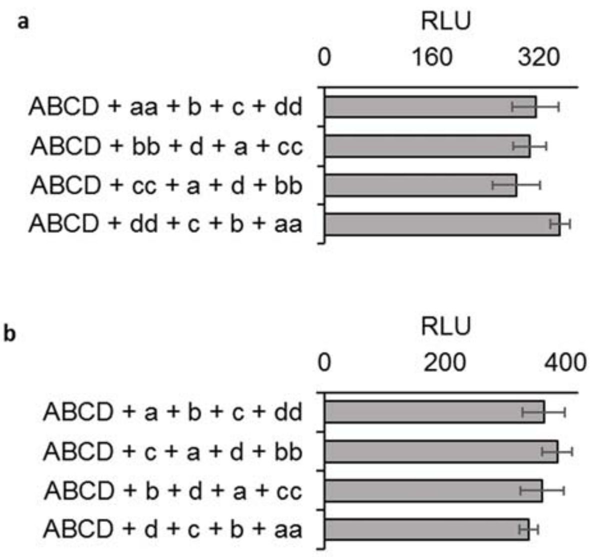


**Figure S11. The identity of DNA tags on the surface of synthetic cells does not influence the fusion events and the signal from the final populations.**
Ancestral synthetic cell population ABCD was mixed with populations containing the same genetic circuit components but decorated with different DNA fusion tags.

### Figure S12

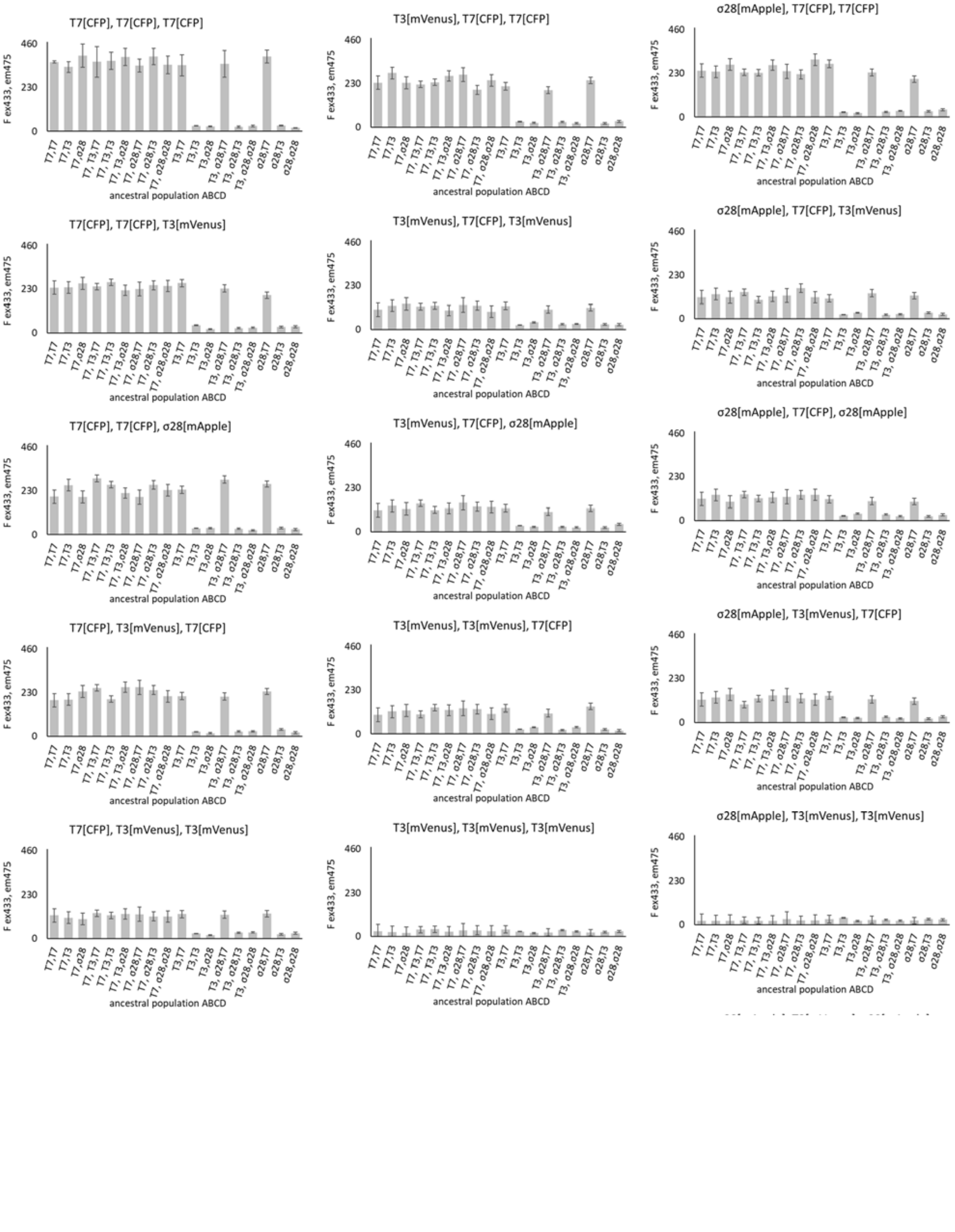
T7[CFP], T7[CFP], T7[CFP]
T3[mVenus], T7[CFP], T7[CFP]
σ28[mApple], T7[CFP], T7[CFP]
T7[CFP], T7[CFP], T3[mVenus]
T3[mVenus], T7[CFP], T3[mVenus]
σ28[mApple], T7[CFP], T3[mVenus]
T7[CFP], T7[CFP], σ28[mApple]
T3[mVenus], T7[CFP], σ28[mApple]
σ28[mApple], T7[CFP], σ28[mApple]
T7[CFP], T3[mVenus], T7[CFP]
T3[mVenus], T3[mVenus], T7[CFP]
σ28[mApple], T3[mVenus], T7[CFP]
T7[CFP], T3[mVenus], T3[mVenus]
T3[mVenus], T3[mVenus], T3[mVenus]
σ28[mApple], T3[mVenus], T3[mVenus]
F ex433, em475
460
230
0
T7,T7
T7,T3
T7,σ28
T7, T3,T7
T7, T3,T3
T7, T3,σ28
T7, σ28,T7
T7, σ28,T3
T7, σ28,σ28
T3,T7
T3,T3
T3,σ28
T3, σ28,T7
T3, σ28,T3
T3, σ28,σ28
σ28,T7
σ28,T3
σ28,σ28
ancestral population ABCD

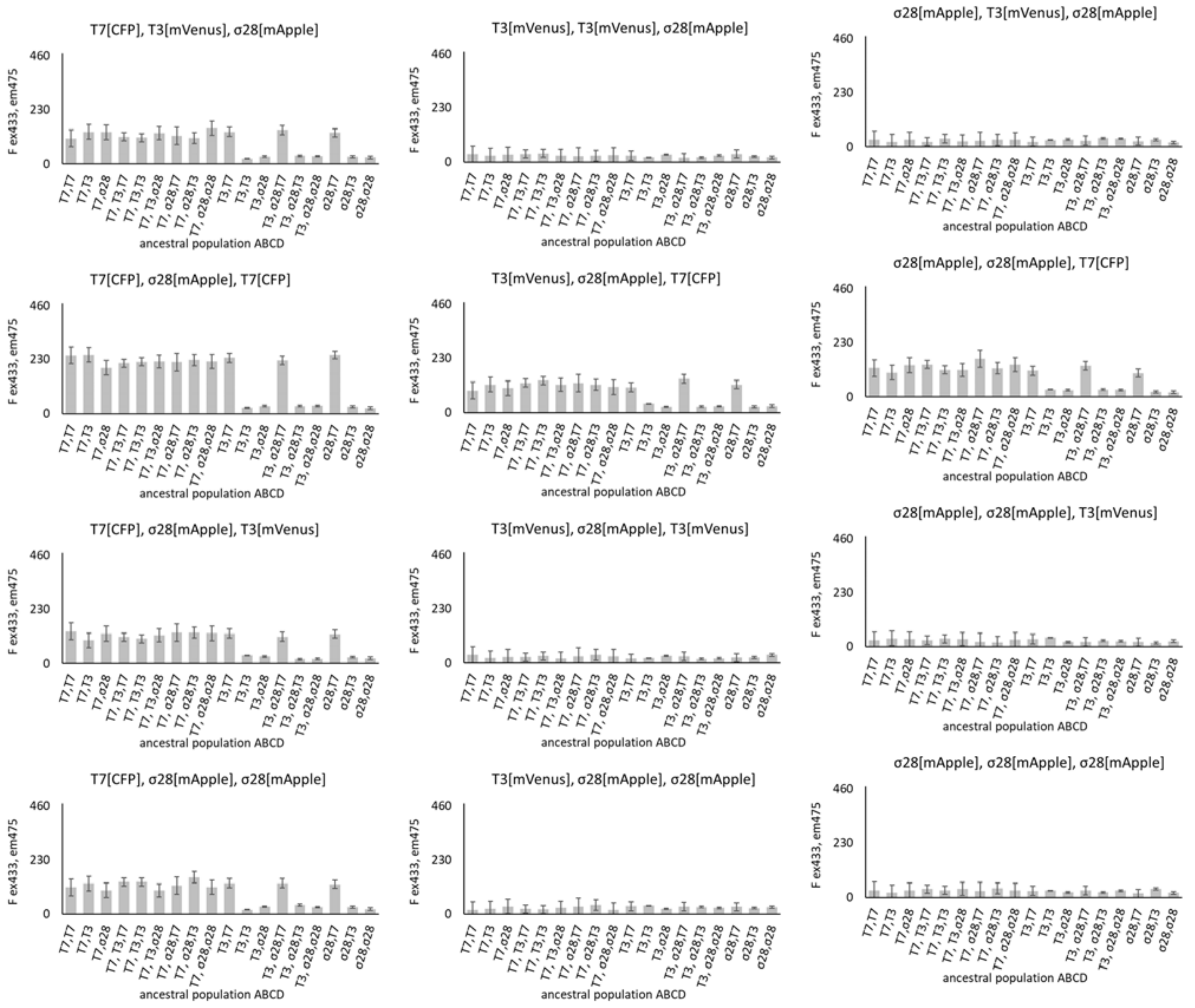


**Figure S12** Activation of independent genetic circuits by subsequent fusion of synthetic cell populations. Here, the output of genetic circuits is measured by the activity of CFP, with excitation wavelength of 433nm and emission wavelength of 475nm. Error bard represent SD for n=3.

### Figure S13

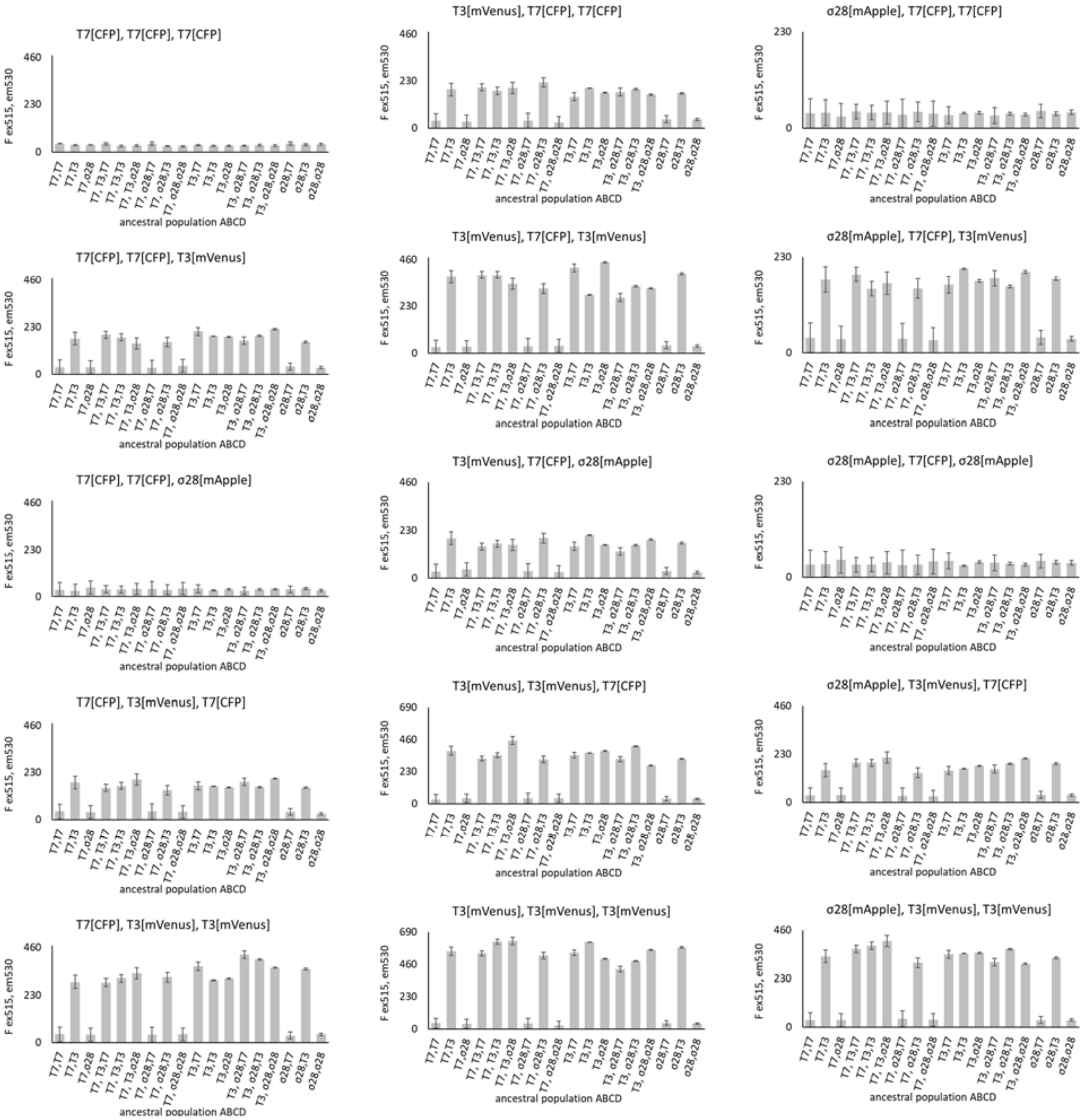

T7[CFP], T7[CFP], T7[CFP]
T3[mVenus], T7[CFP], T7[CFP]
σ28[mApple], T7[CFP], T7[CFP]
T7[CFP], T7[CFP], T3[mVenus]
T3[mVenus], T7[CFP], T3[mVenus]
σ28[mApple], T7[CFP], T3[mVenus]
T7[CFP], T7[CFP], σ28[mApple]
T3[mVenus], T7[CFP], σ28[mApple]
σ28[mApple], T7[CFP], σ28[mApple]
T7[CFP], T3[mVenus], T7[CFP]
T3[mVenus], T3[mVenus], T7[CFP]
σ28[mApple], T3[mVenus], T7[CFP]
T7[CFP], T3[mVenus], T3[mVenus]
T3[mVenus], T3[mVenus], T3[mVenus]
σ28[mApple], T3[mVenus], T3[mVenus]
F ex515, em530
ancestral population ABCD
0
230
460
690

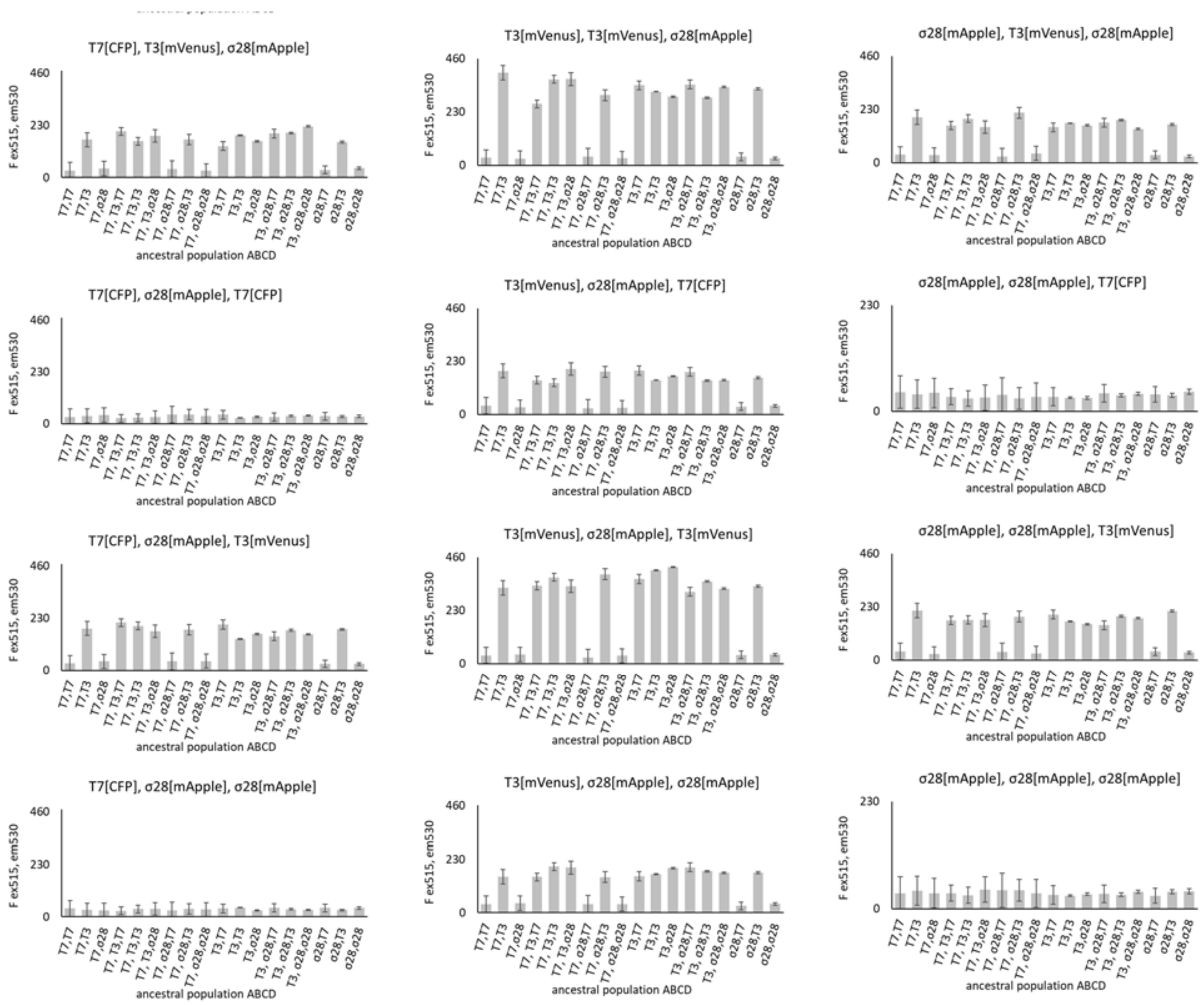


**Figure S13** Activation of independent genetic circuits by subsequent fusion of synthetic cell populations. Here, the output of genetic circuits is measured by the activity of mVenus reporter protein, with excitation wavelength of 515nm and emission wavelength of 530nm. Error bard represent SD for n=3.

### Figure S14

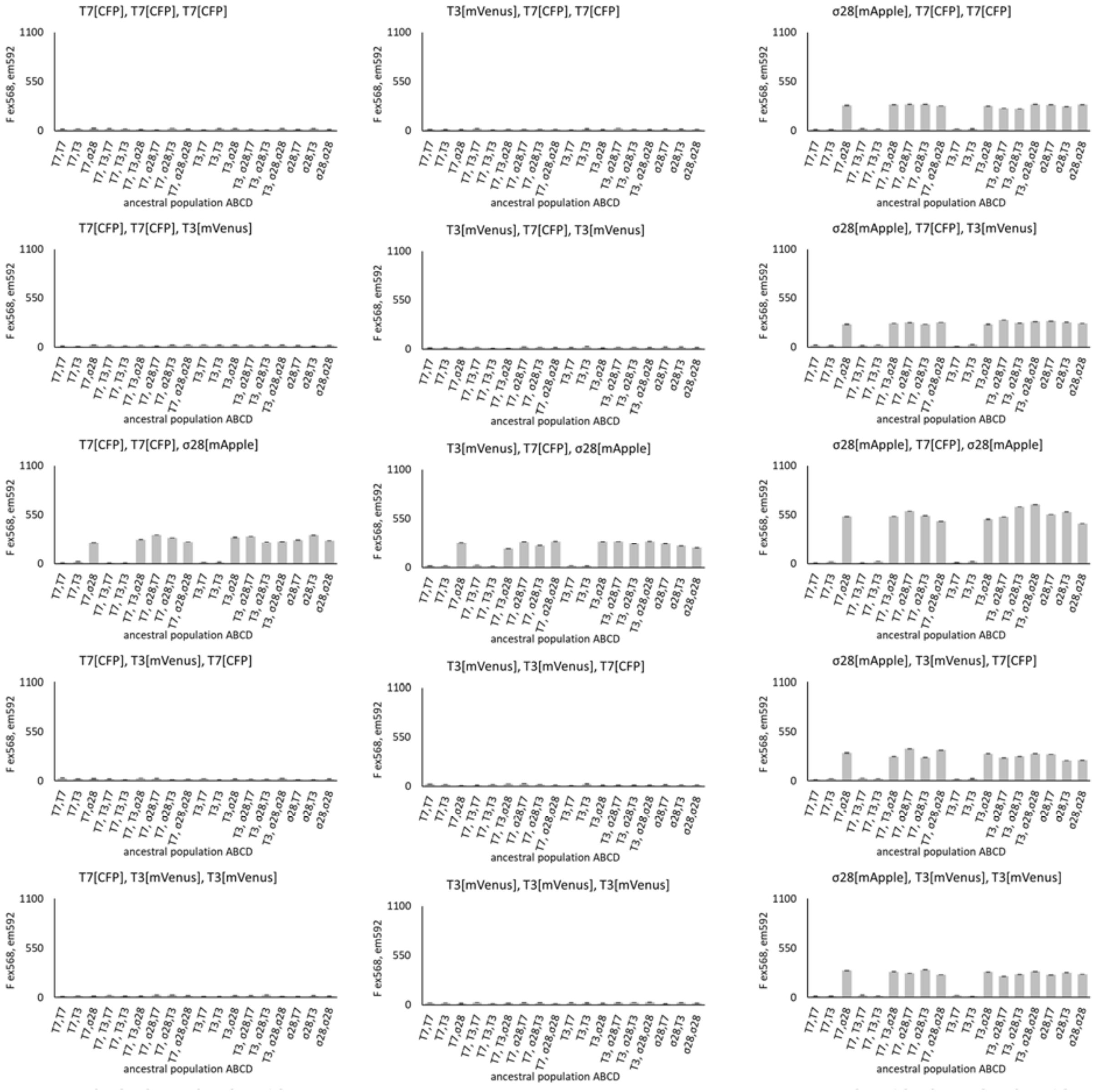
T7[CFP], T7[CFP], T7[CFP]
T3[mVenus], T7[CFP], T7[CFP]
σ28[mApple], T7[CFP], T7[CFP]
T7[CFP], T7[CFP], T3[mVenus]
T3[mVenus], T7[CFP], T3[mVenus]
σ28[mApple], T7[CFP], T3[mVenus]
T7[CFP], T7[CFP], σ28[mApple]
T3[mVenus], T7[CFP], σ28[mApple]
σ28[mApple], T7[CFP], σ28[mApple]
T7[CFP], T3[mVenus], T7[CFP]
T3[mVenus], T3[mVenus], T7[CFP]
σ28[mApple], T3[mVenus], T7[CFP]
T7[CFP], T3[mVenus], T3[mVenus]
T3[mVenus], T3[mVenus], T3[mVenus]
σ28[mApple], T3[mVenus], T3[mVenus]
F ex568, em592
1100
550
0
T7,T7
T7,T3
T7,σ28
T7, T3,T7
T7, T3,T3
T7, T3,σ28
T7, σ28,T7
T7, σ28,T3
T7, σ28,σ28
T3,T7
T3,T3
T3,σ28
T3, σ28,T7
T3, σ28,T3
T3, σ28,σ28
σ28,T7
σ28,T3
σ28,σ28
ancestral population ABCD

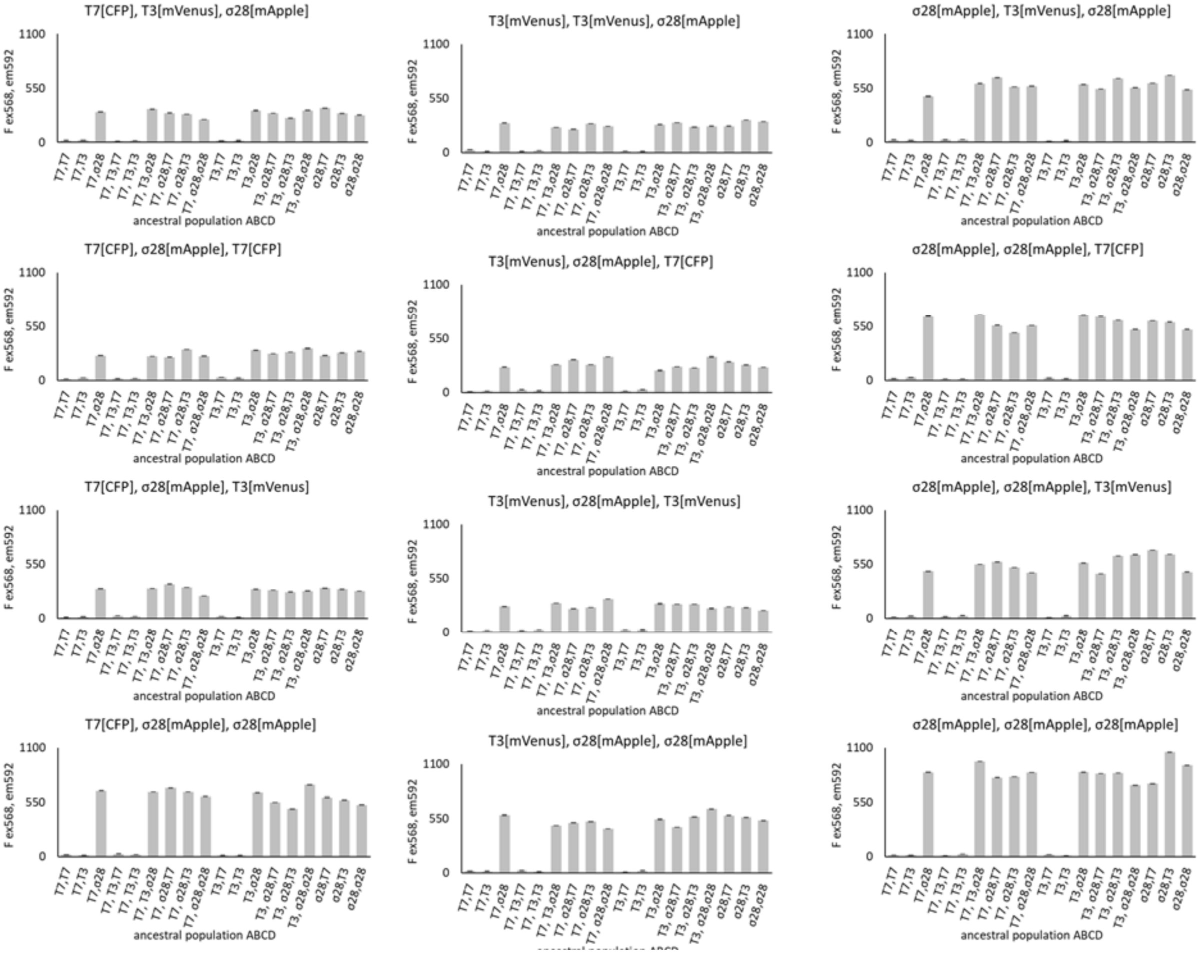


**Figure S14** Activation of independent genetic circuits by subsequent fusion of synthetic cell populations. Here, the output of genetic circuits is measured by the activity of mApple reporter protein, with excitation wavelength of 568nm and emission wavelength of 592nm. Error bard represent SD for n=3.

### Figure S15

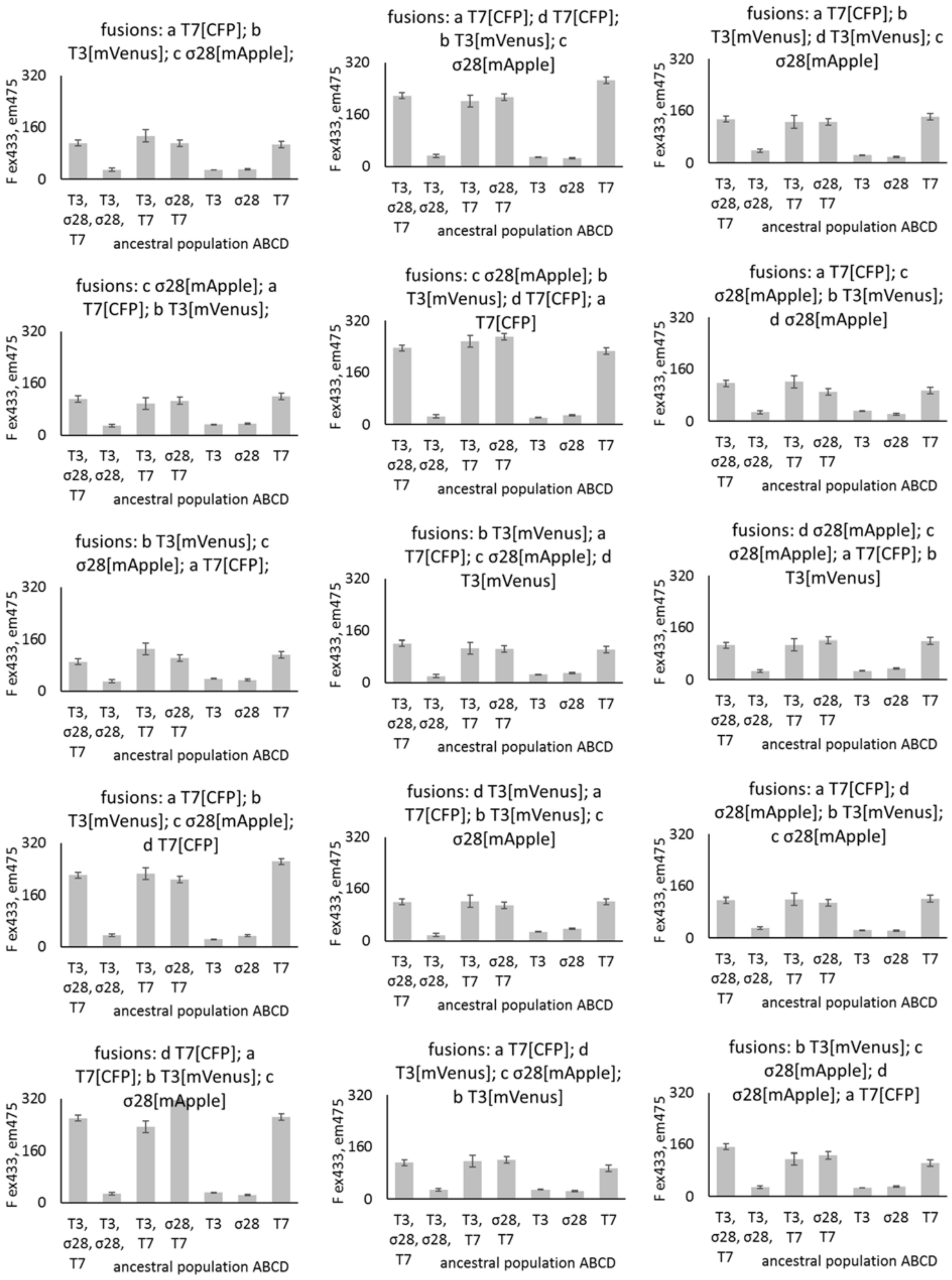
fusions: a T7[CFP]; b T3[mVenus]; c σ28[mApple];
fusions: a T7[CFP]; d T7[CFP]; b T3[mVenus]; c σ28[mApple]
fusions: a T7[CFP]; b T3[mVenus]; d T3[mVenus]; c σ28[mApple]
fusions: c σ28[mApple]; a T7[CFP]; b T3[mVenus];
fusions: c σ28[mApple]; b T3[mVenus]; d T7[CFP]; a T7[CFP]
fusions: a T7[CFP]; c σ28[mApple]; b T3[mVenus]; d σ28[mApple]
fusions: b T3[mVenus]; c σ28[mApple]; a T7[CFP];
fusions: b T3[mVenus]; a T7[CFP]; c σ28[mApple]; d T3[mVenus]
fusions: d σ28[mApple]; c σ28[mApple]; a T7[CFP]; b T3[mVenus]
fusions: a T7[CFP]; b T3[mVenus]; c σ28[mApple]; d T7[CFP]
fusions: d T3[mVenus]; a T7[CFP]; b T3[mVenus]; c σ28[mApple]
fusions: a T7[CFP]; d σ28[mApple]; b T3[mVenus]; c σ28[mApple]
fusions: d T7[CFP]; a T7[CFP]; b T3[mVenus]; c σ28[mApple]
fusions: a T7[CFP]; d T3[mVenus]; c σ28[mApple]; b T3[mVenus]
fusions: b T3[mVenus]; c σ28[mApple]; d σ28[mApple]; a T7[CFP]
F ex433, em475
0
160
320
T3, σ28, T7
T3, σ28,
T3, T7
σ28, T7
T3
σ28
T7
ancestral population ABCD

**Figure S15. Order of population fusions does not affect the final signal measured in the final descendant population of synthetic cells.** Here, the output of genetic circuits is measured by the activity of CFP, with excitation wavelength of 433nm and emission wavelength of 475nm. Error bard represent SD for n=3.

### Figure S16

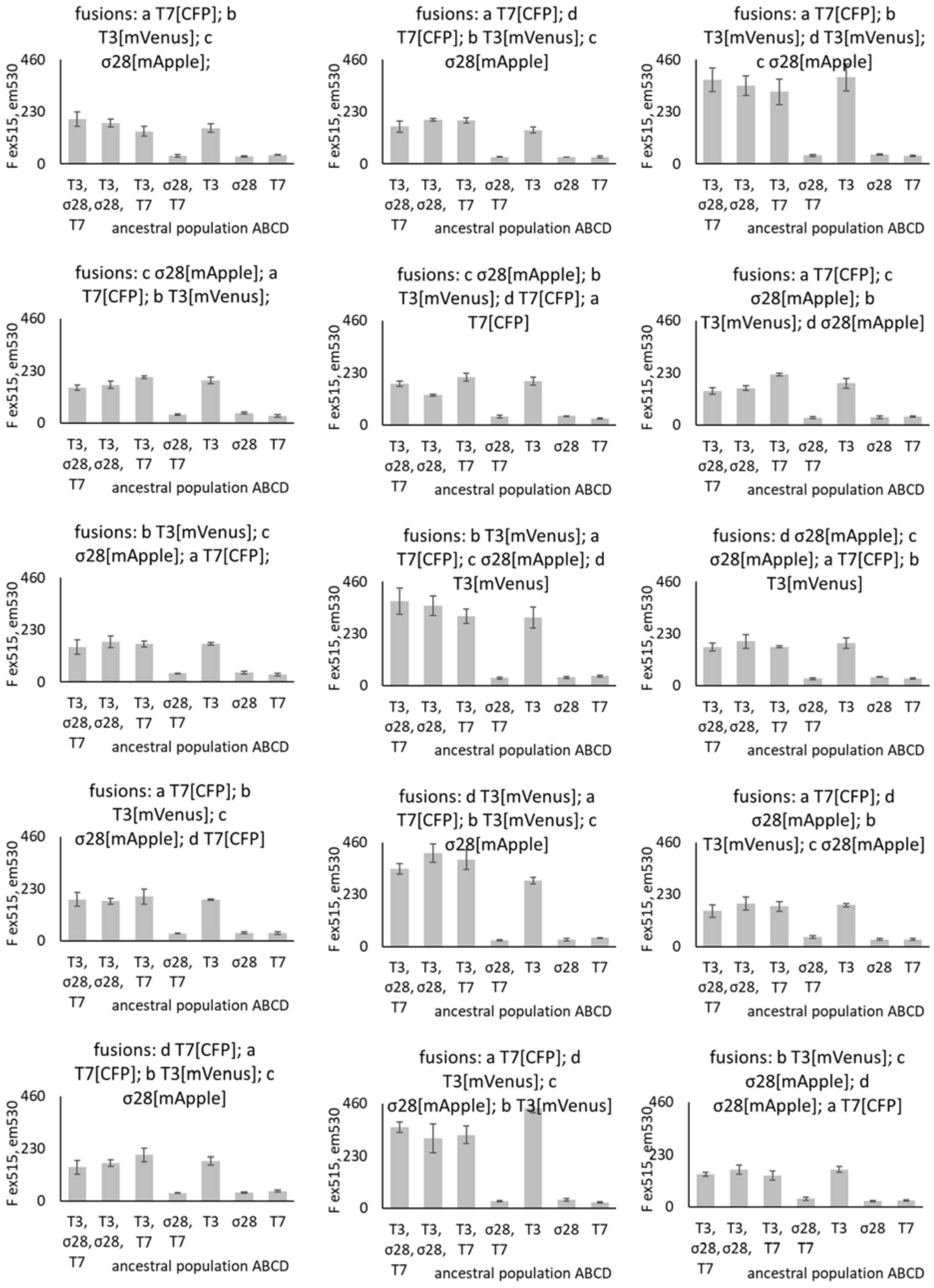

fusions: a T7[CFP]; b T3[mVenus]; c σ28[mApple];
fusions: a T7[CFP]; d T7[CFP]; b T3[mVenus]; c σ28[mApple]
fusions: a T7[CFP]; b T3[mVenus]; d T3[mVenus]; c σ28[mApple]
fusions: c σ28[mApple]; a T7[CFP]; b T3[mVenus];
fusions: c σ28[mApple]; b T3[mVenus]; d T7[CFP]; a T7[CFP]
fusions: a T7[CFP]; c σ28[mApple]; b T3[mVenus]; d σ28[mApple]
fusions: b T3[mVenus]; c σ28[mApple]; a T7[CFP];
fusions: b T3[mVenus]; a T7[CFP]; c σ28[mApple]; d T3[mVenus]
fusions: d σ28[mApple]; c σ28[mApple]; a T7[CFP]; b T3[mVenus]
fusions: a T7[CFP]; b T3[mVenus]; c σ28[mApple]; d T7[CFP]
fusions: d T3[mVenus]; a T7[CFP]; b T3[mVenus]; c σ28[mApple]
fusions: a T7[CFP]; d σ28[mApple]; b T3[mVenus]; c σ28[mApple]
fusions: d T7[CFP]; a T7[CFP]; b T3[mVenus]; c σ28[mApple]
fusions: a T7[CFP]; d T3[mVenus]; c σ28[mApple]; b T3[mVenus]
fusions: b T3[mVenus]; c σ28[mApple]; d σ28[mApple]; a T7[CFP]
F ex515, em530
0
230
460
T3, σ28, T7
T3, σ28,
T3, T7
σ28, T7
T3
σ28
T7
ancestral population ABCD

**Figure S16. Order of population fusions does not affect the final signal measured in the final descendant population of synthetic cells.** Here, the output of genetic circuits is measured by the activity of mVenus reporter protein, with excitation wavelength of 515nm and emission wavelength of 530nm. Error bard represent SD for n=3.

### Figure S17

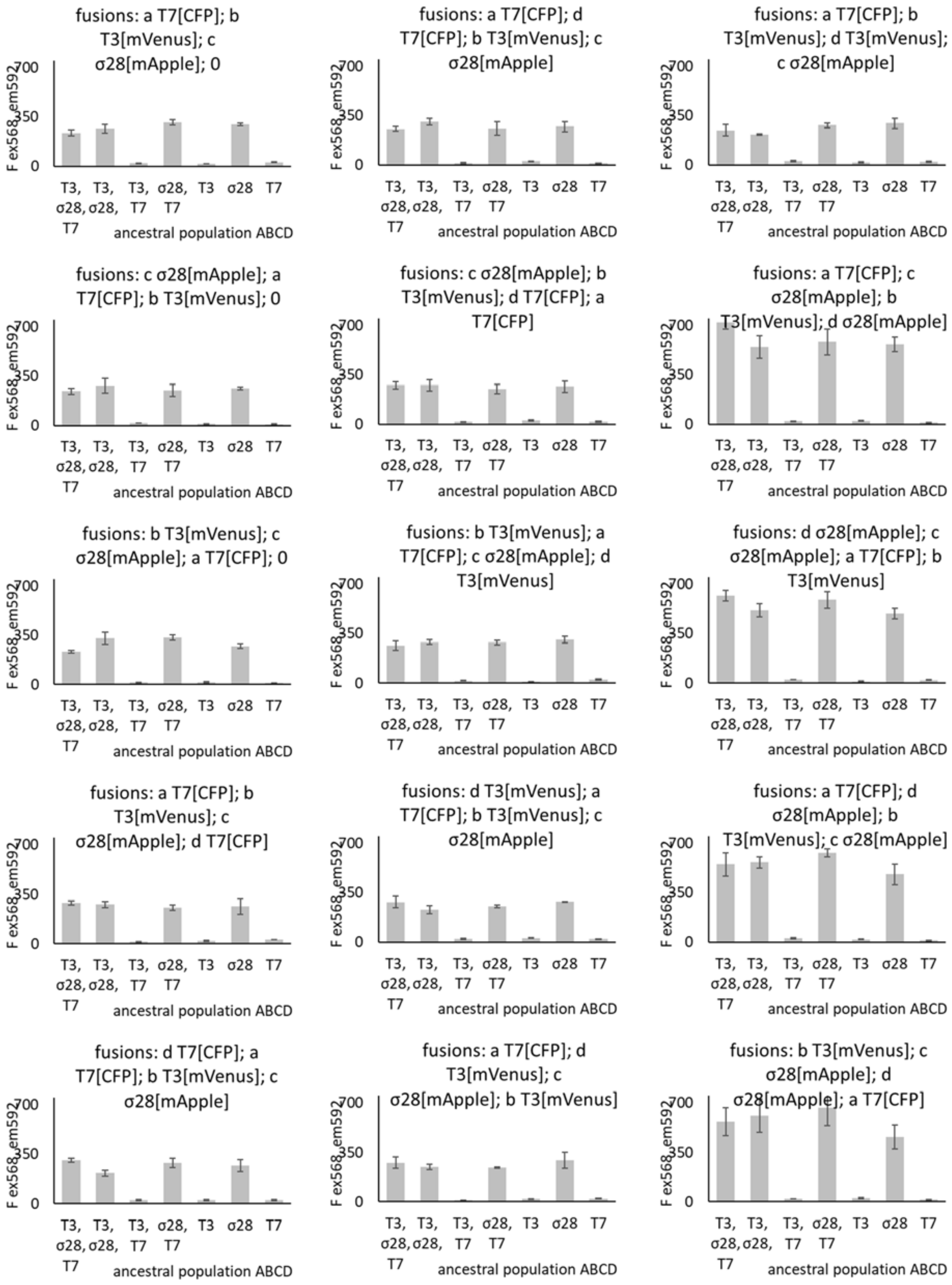

fusions: a T7[CFP]; b T3[mVenus]; c σ28[mApple]; 0
fusions: a T7[CFP]; d T7[CFP]; b T3[mVenus]; c σ28[mApple]
fusions: a T7[CFP]; b T3[mVenus]; d T3[mVenus]; c σ28[mApple]
fusions: c σ28[mApple]; a T7[CFP]; b T3[mVenus]; 0
fusions: c σ28[mApple]; b T3[mVenus]; d T7[CFP]; a T7[CFP]
fusions: a T7[CFP]; c σ28[mApple]; b T3[mVenus]; d σ28[mApple]
fusions: b T3[mVenus]; c σ28[mApple]; a T7[CFP]; 0
fusions: b T3[mVenus]; a T7[CFP]; c σ28[mApple]; d T3[mVenus]
fusions: d σ28[mApple]; c σ28[mApple]; a T7[CFP]; b T3[mVenus]
fusions: a T7[CFP]; b T3[mVenus]; c σ28[mApple]; d T7[CFP]
fusions: d T3[mVenus]; a T7[CFP]; b T3[mVenus]; c σ28[mApple]
fusions: a T7[CFP]; d σ28[mApple]; b T3[mVenus]; c σ28[mApple]
fusions: d T7[CFP]; a T7[CFP]; b T3[mVenus]; c σ28[mApple]
fusions: a T7[CFP]; d T3[mVenus]; c σ28[mApple]; b T3[mVenus]
fusions: b T3[mVenus]; c σ28[mApple]; d σ28[mApple]; a T7[CFP]
F ex568, em592
700
350
0
T3, σ28, T7
T3, σ28,
T3, T7
σ28, T7
T3
σ28
T7
ancestral population ABCD

**Figure S17. Order of population fusions does not affect the final signal measured in the final descendant population of synthetic cells.** Here, the output of genetic circuits is measured by the activity of mApple reporter protein, with excitation wavelength of 568nm and emission wavelength of 592nm. Error bard represent SD for n=3.

### Figure S18

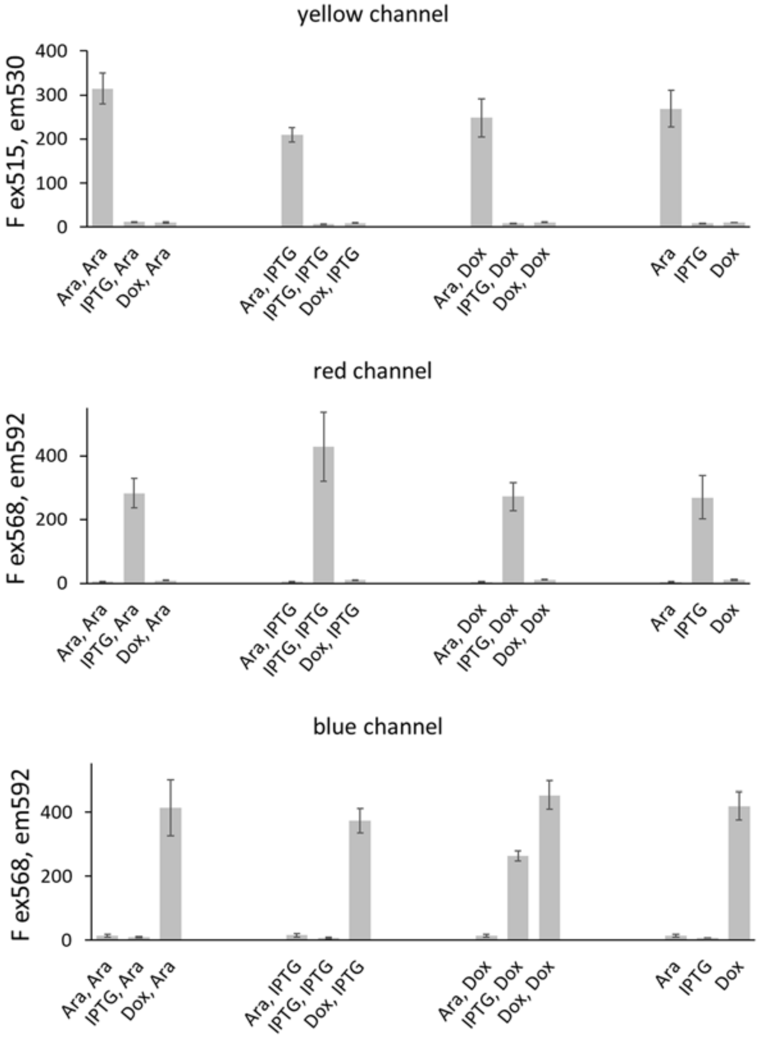


**Figure S18** Individual data points for small molecule inducible differentiation of synthetic cell populations.
Yellow mVenus channel ex 515nm and em 530nm, red mApple channel ex 568nm and em 592nm, blue CFP channel ex 433nm and em 475nm. Each sample is arithmetic averae of three replicates, error bars are S.E.M. for n=3.

### Figure S19

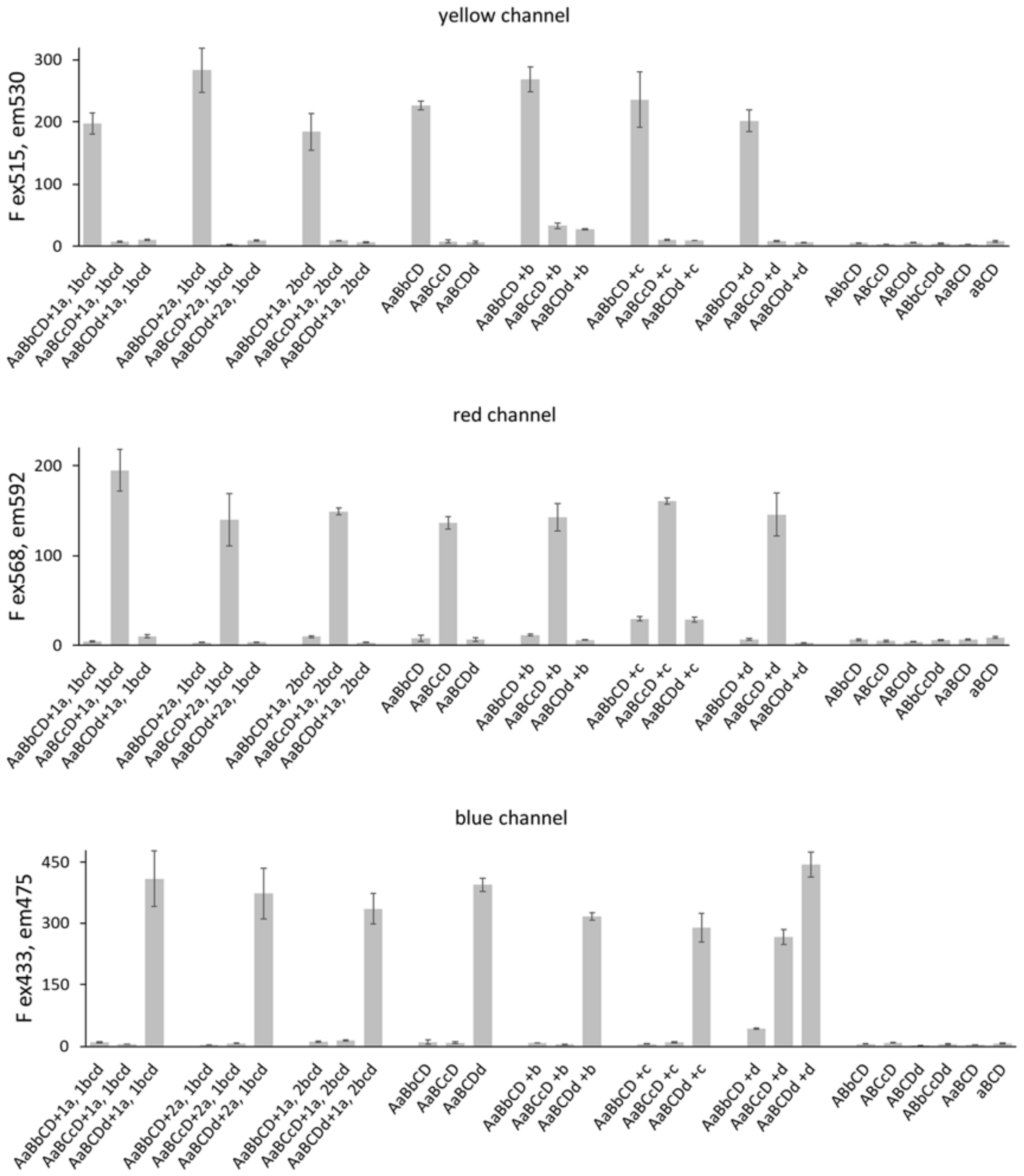


**Figure S19** Data points for fluorescent protein measurements in synthetic cells differentiated from ancestral pluripotent population.
Yellow mVenus channel ex 515nm and em 530nm, red mApple channel ex 568nm and em 592nm, blue CFP channel ex 433nm and em 475nm. Each sample is arithmetic averae of three replicates, error bars are S.E.M. for n=3.

### Figure S20

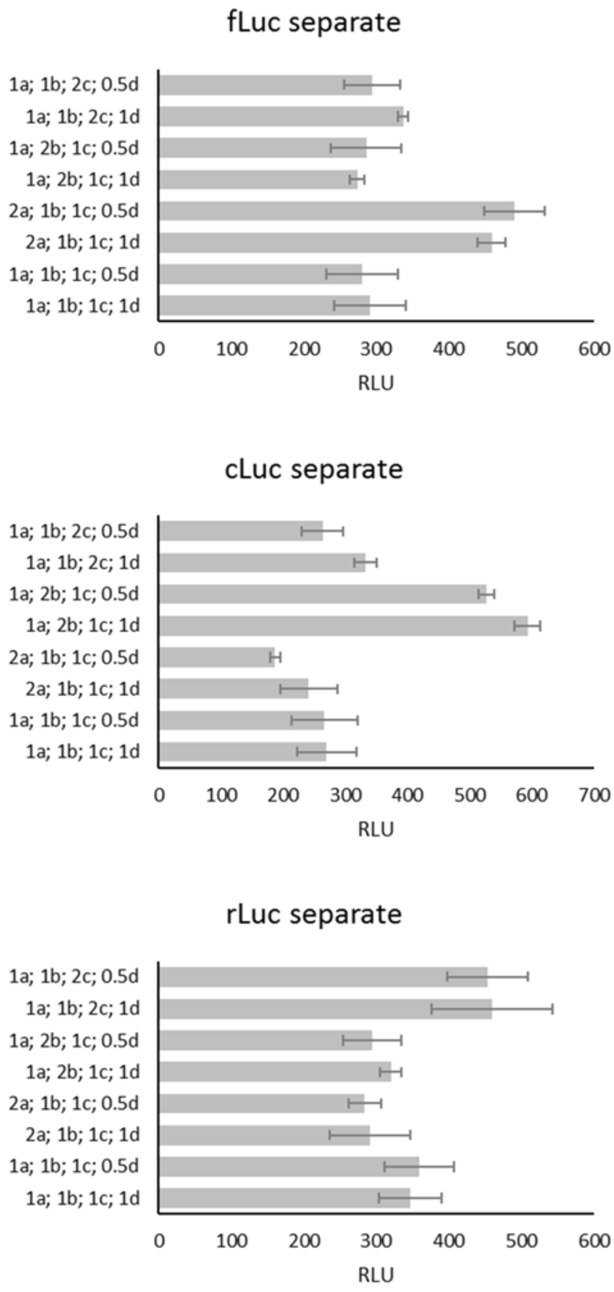


**Figure S20** Differentiation experiments carried with the lineages being physically separated in different tubes. Error bars represent +/- standard deviation, n=3.

### Figure S21

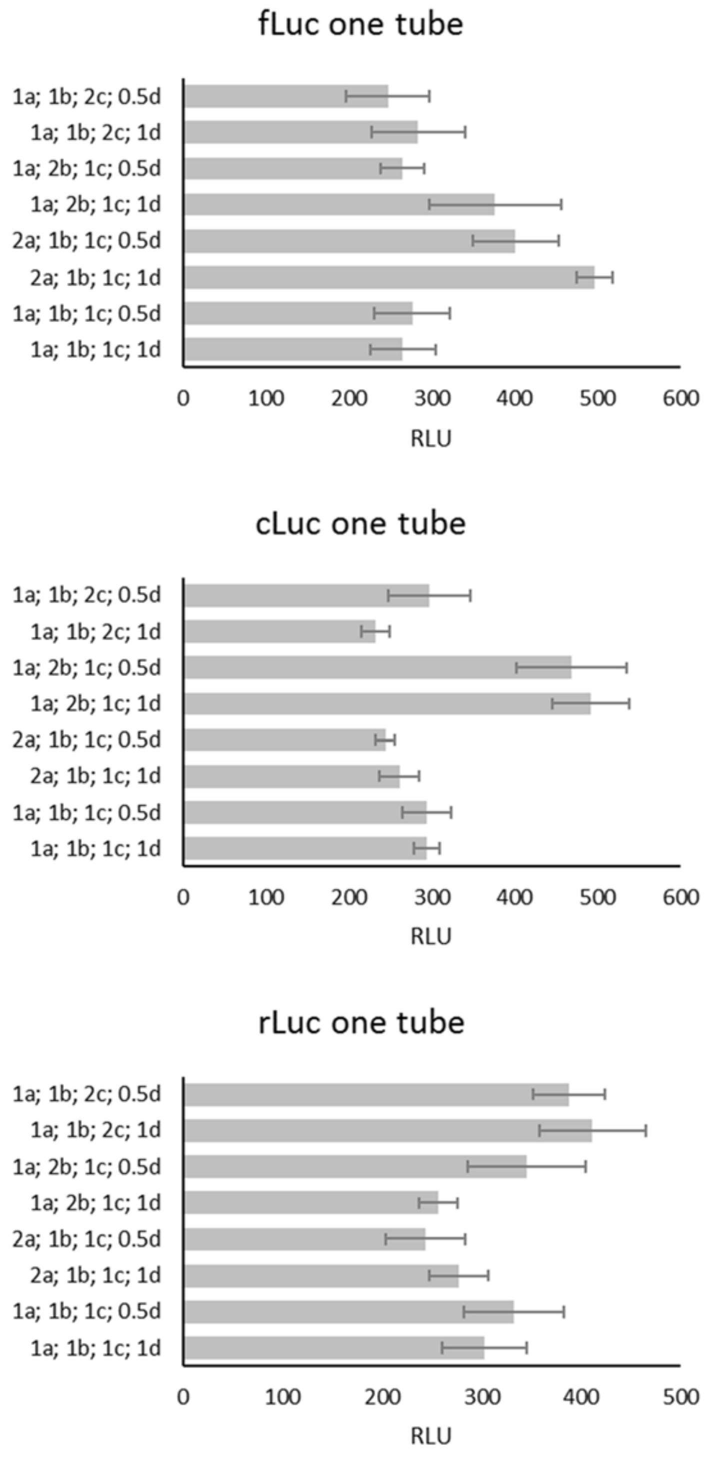


**Figure S21** Differentiation experiments carried with the lineages incubated together in the same tube. Error bars represent +/- standard deviation, n=3.

### Figure S22

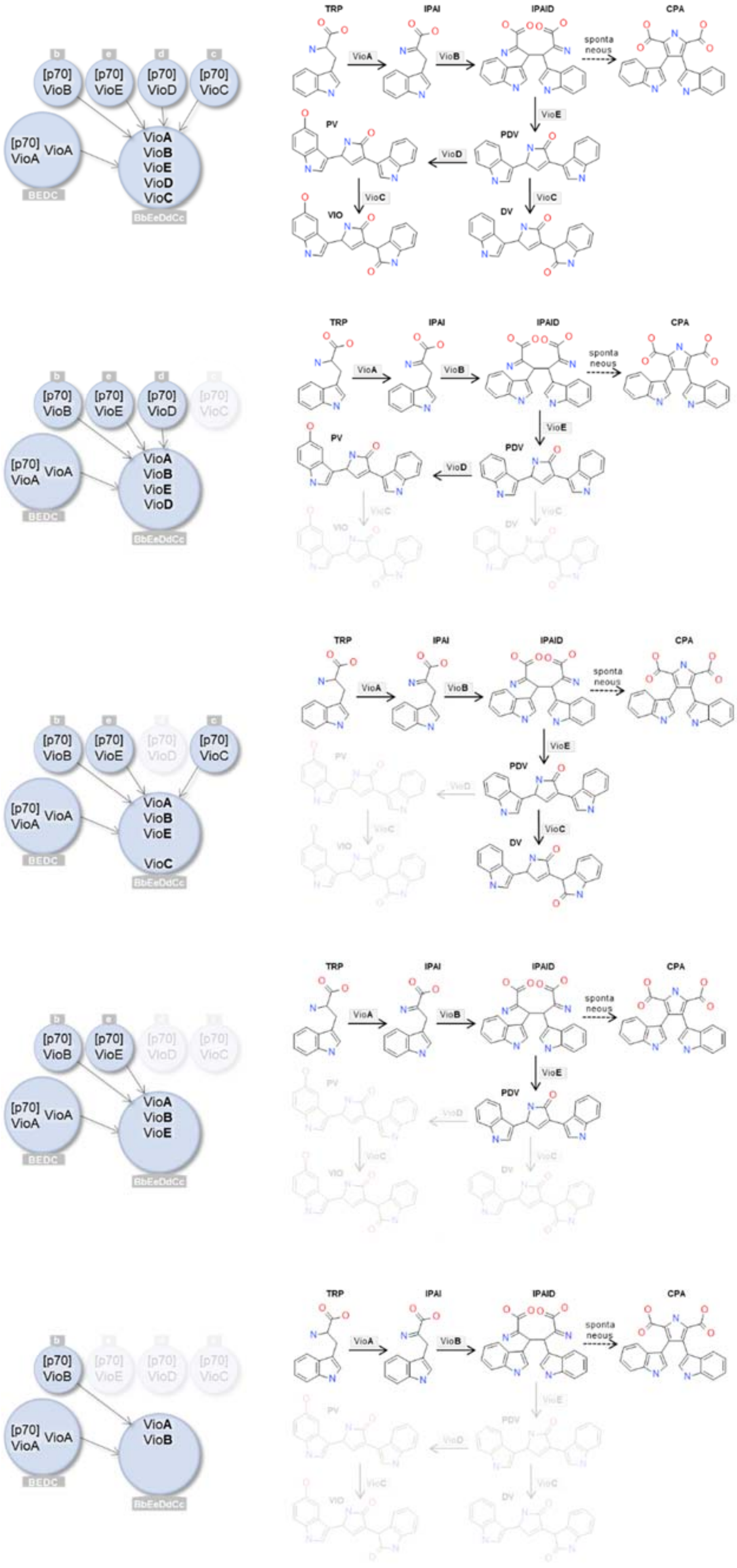


**Figure S22** Variants of the Vio pathway, built by different series of synthetic cell populations carrying Vio enzymes.

### Figure S23

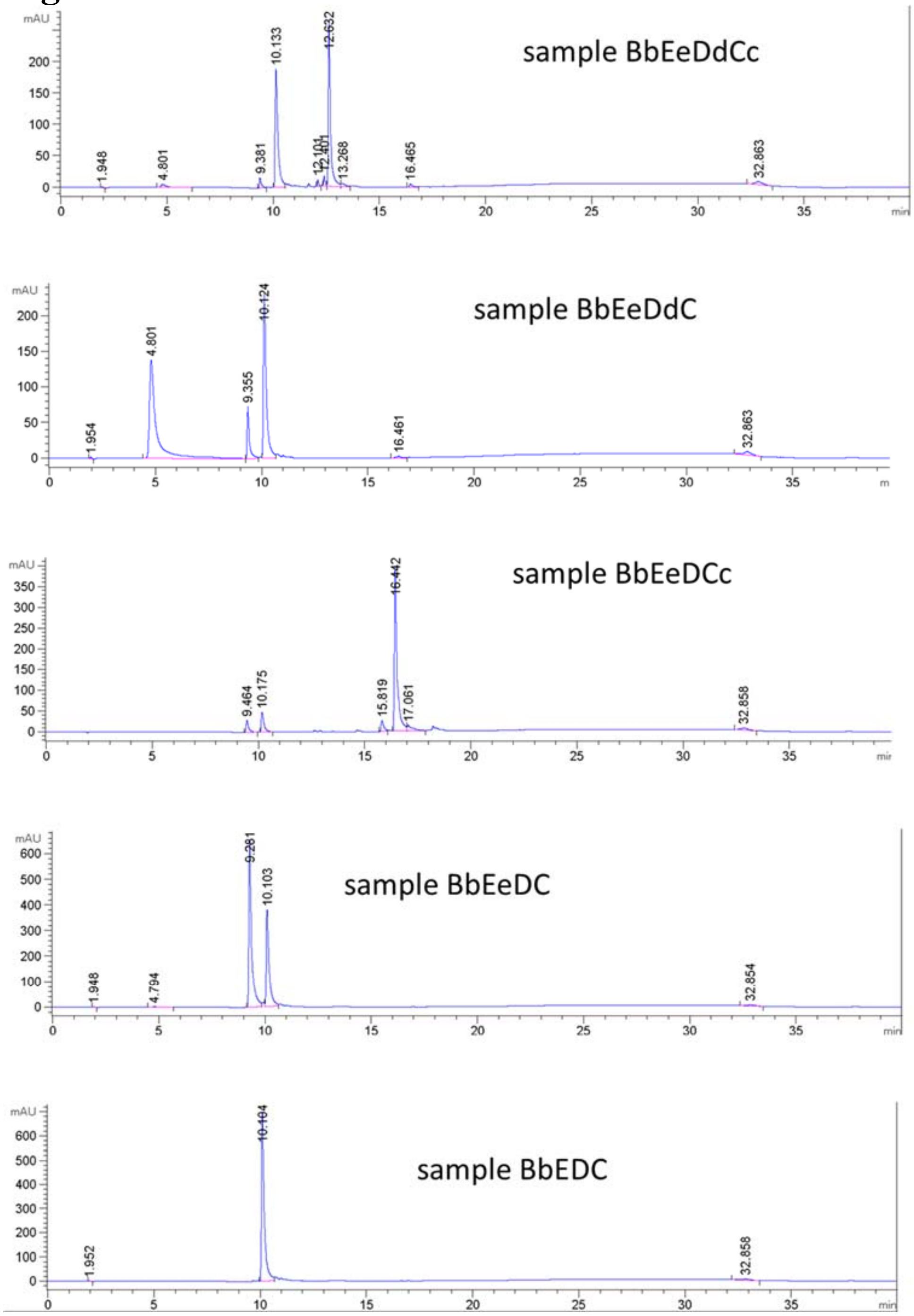


**Figure S23**. HPLC traces of violacein pathway samples. Absorbance measured at 570nm. Each chromatogram represents one of three replicates of each experiment.

### Figure S24

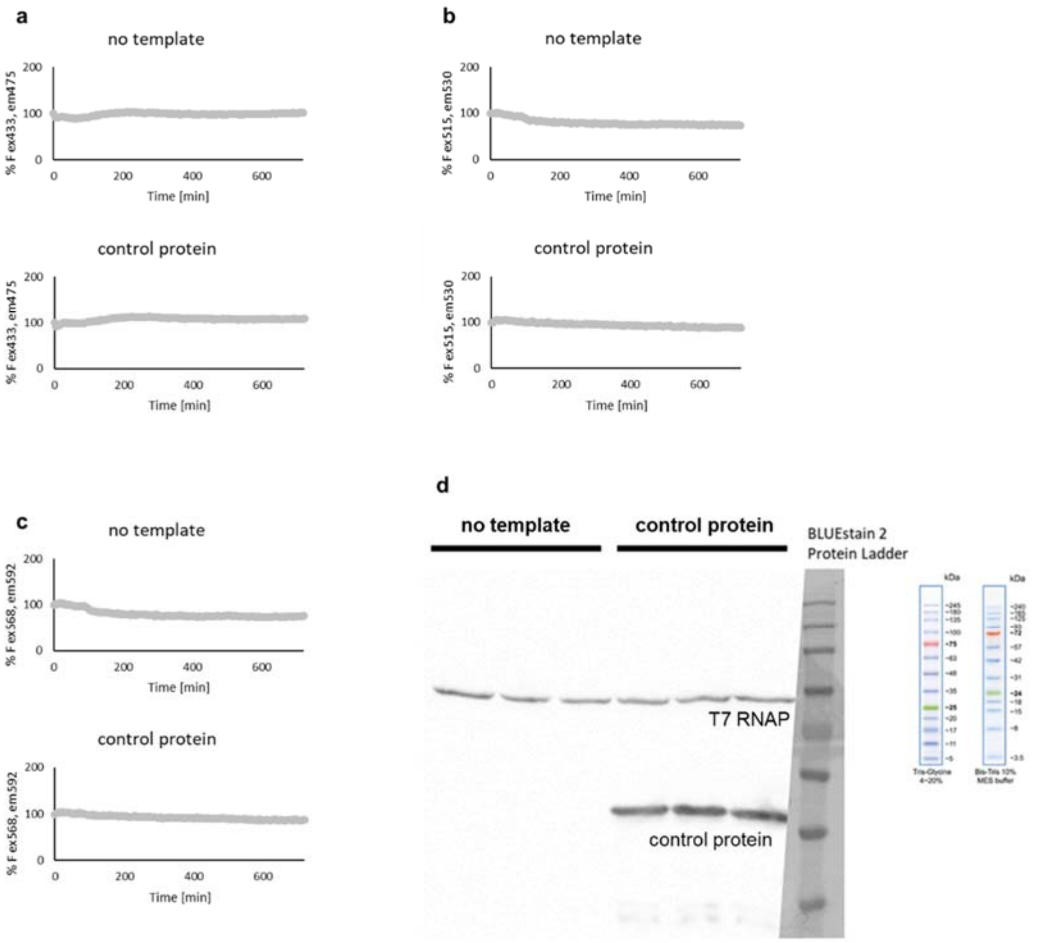


**Figure S24** Fluorescence background does not change with incubation of TxTl mixture over time, in presence and absence of translation.

Samples were incubated with and without plasmid DNA, and fluorescence was monitored on three channels: **a** blue CFP channel $\lambda_{ex}$ 433nm and $\lambda_{em}$ 475nm, **b**: yellow mVenus channel $\lambda_{ex}$ 515nm and $\lambda_{em}$ 530nm, **c**: red mApple channel $\lambda_{ex}$ 568nm and $\lambda_{em}$ 592nm.Each trace is arithmetic average of three replicates.

**d**: Western blot analysis of all samples (2 samples: without DNA gene and with control protein plasmid, each in triplicate), with 6xHis antibody.

The control protein is OphA protein from *Omphalotus olearius* Jack-o'-Lantern mushroom, a periplasmic ligand binding protein chosen here as an example of non-fluorescent protein that robustly expresses in bacterial TxTl.

### Figure S25

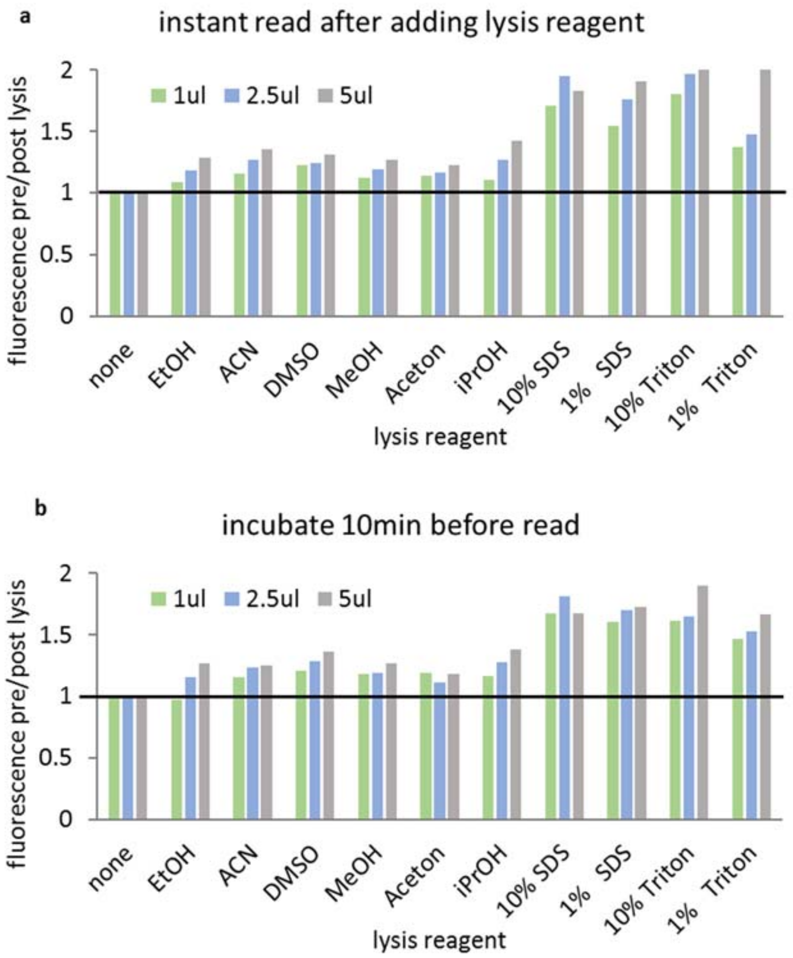


**Figure S25. Testing different lysis conditions for liposomes.**

Liposome samples were prepared with 30mM calcein, purified on size exclusion column to remove unencapsulated dye, liposome fractions were pooled into one. Liposome solution was divided into 50uL aliquots, fluorescence was measured, and 1uL, 2.5uL or 5uL of stock lysis solution was added. Fluorescence was measured again immediately after adding lysis solution and then again after 10min incubation at room temperature.

The lysis efficiency, measured as increased signal of calcein as it dequenches upon release from lysed liposomes, is reported as ratio of fluorescence before and after lysis. The black horizontal line shows fluorescence ratio in the absence of lysis, when no change of fluorescence is observed.

Experiment conditions: 25 mM tris-HCl pH 7.5, 5mM liposomes. Reported values are arithmetical average of three replicates.

### Figure S26

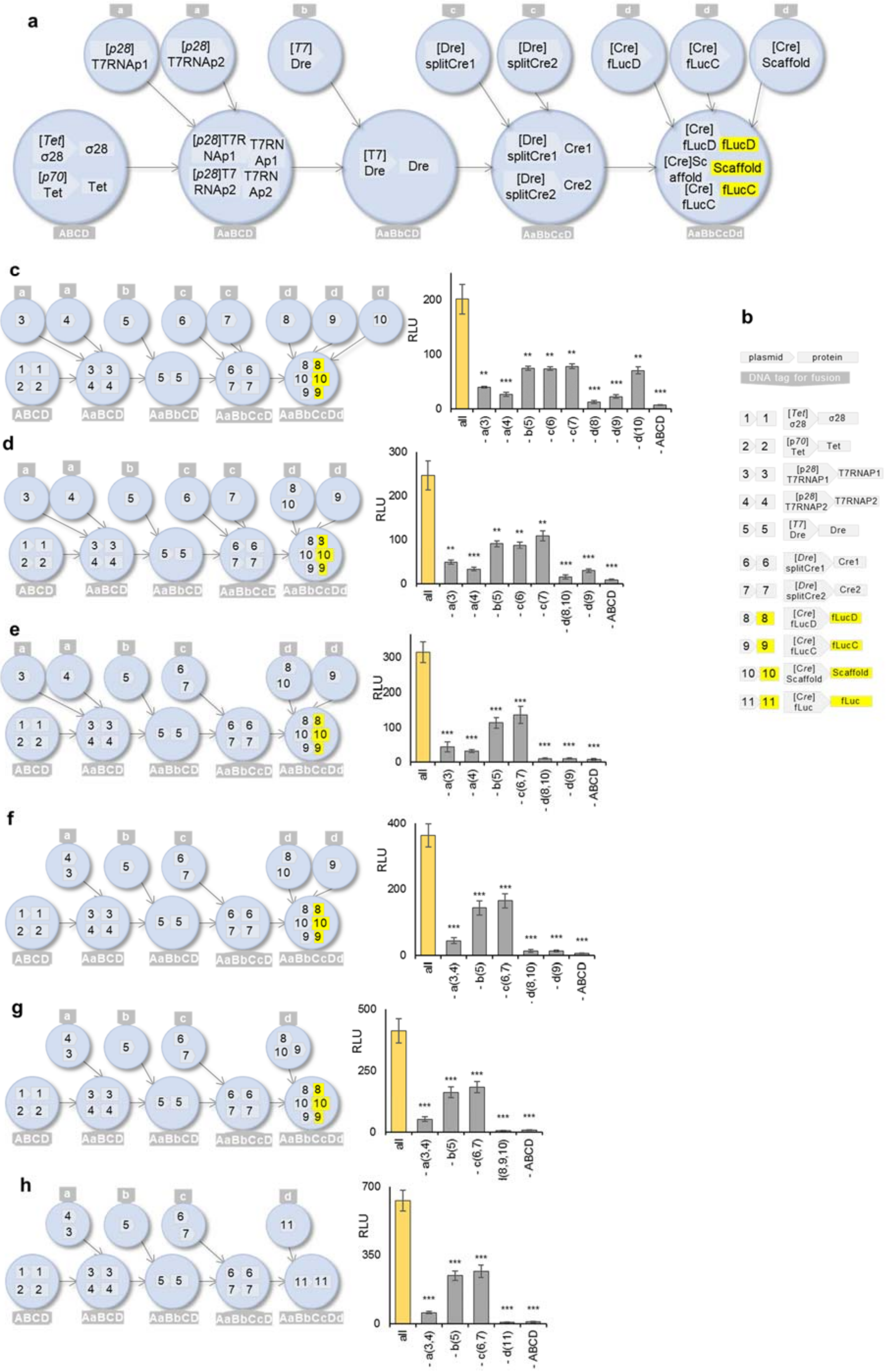

a
[p28] T7RNAp1
[p28] T7RNAp2
[T7] Dre
[Dre] splitCre1
[Dre] splitCre2
[Cre] fLucD
[Cre] fLucC
[Cre] Scaffold
[Tet] σ28
σ28
[p70] Tet
Tet
ABCD
AaBCD
AaBbCD
AaBbCcD
AaBbCcDd
b
plasmid
protein
DNA tag for fusion
c
d
e
f
g
h
RLU
all
- a(3)
- a(4)
- b(5)
- c(6)
- c(7)
- d(8)
- d(9)
- d(10)
- ABCD
- d(8,10)
- c(6,7)
- a(3,4)
- d(11)

**Figure S26** Fusion of synthetic minimal cells containing genetic circuits creates linear (a-h) combinatorial pathways.
**a**: Linear pathway is constructed with series of RNA polymerases and recombinases, encapsulated in different populations of synthetic minimal cells. Sequential fusion of populations creates the final circuit; readout of the pathway is activity of firefly luciferase reconstituted from three-partite split reporter. The detailed schematic of the pathway is on **Figure S6**.
**b**: symbols used on panel a, with corresponding shorthand numbers used in panels c-h. Arrows indicate promoters, squares indicate protein products.
**c**-**h**: Linear genetic pathways of varying complexity. Synthetic cells with membrane decorated with DNA tags were mixed with ancestral synthetic cell population ABCD. The content of the fusing liposomes, not the specific identity of the membrane tags, influences the final population state (**Fig. S11**). Variants of those pathways were first tested via simultaneous expression in one solution, without encapsulation in synthetic cells (**Fig. S7**).

#### Figure S27

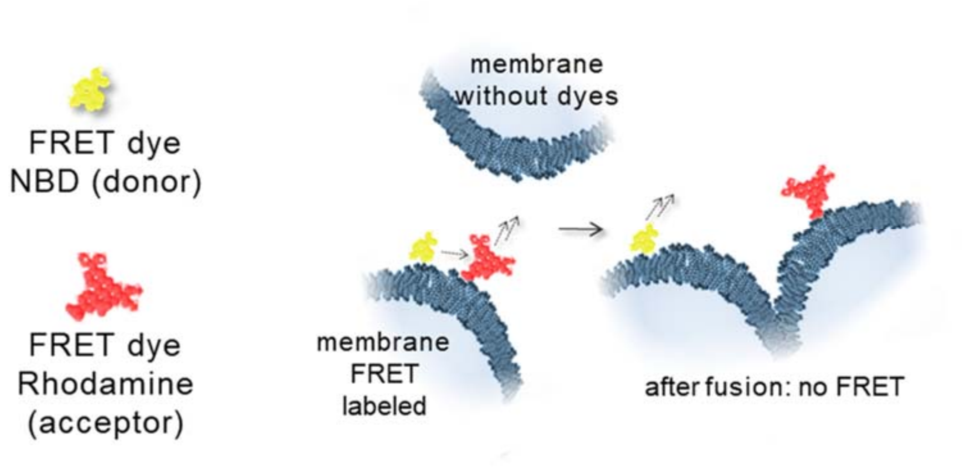


**Figure S27** The schematic of the FRET membrane mixing assay with membranes labeled with NBD-PE (N-(7-Nitrobenz-2-Oxa-1,3-Diazol-4-yl)-1,2-Dihexadecanoyl-sn-Glycero-3-Phosphoethanolamine) and Lissamine Rhodamine B (1,2-Dihexadecanoyl-sn-Glycero-3-Phosphoethanolamine).

### Figure S28

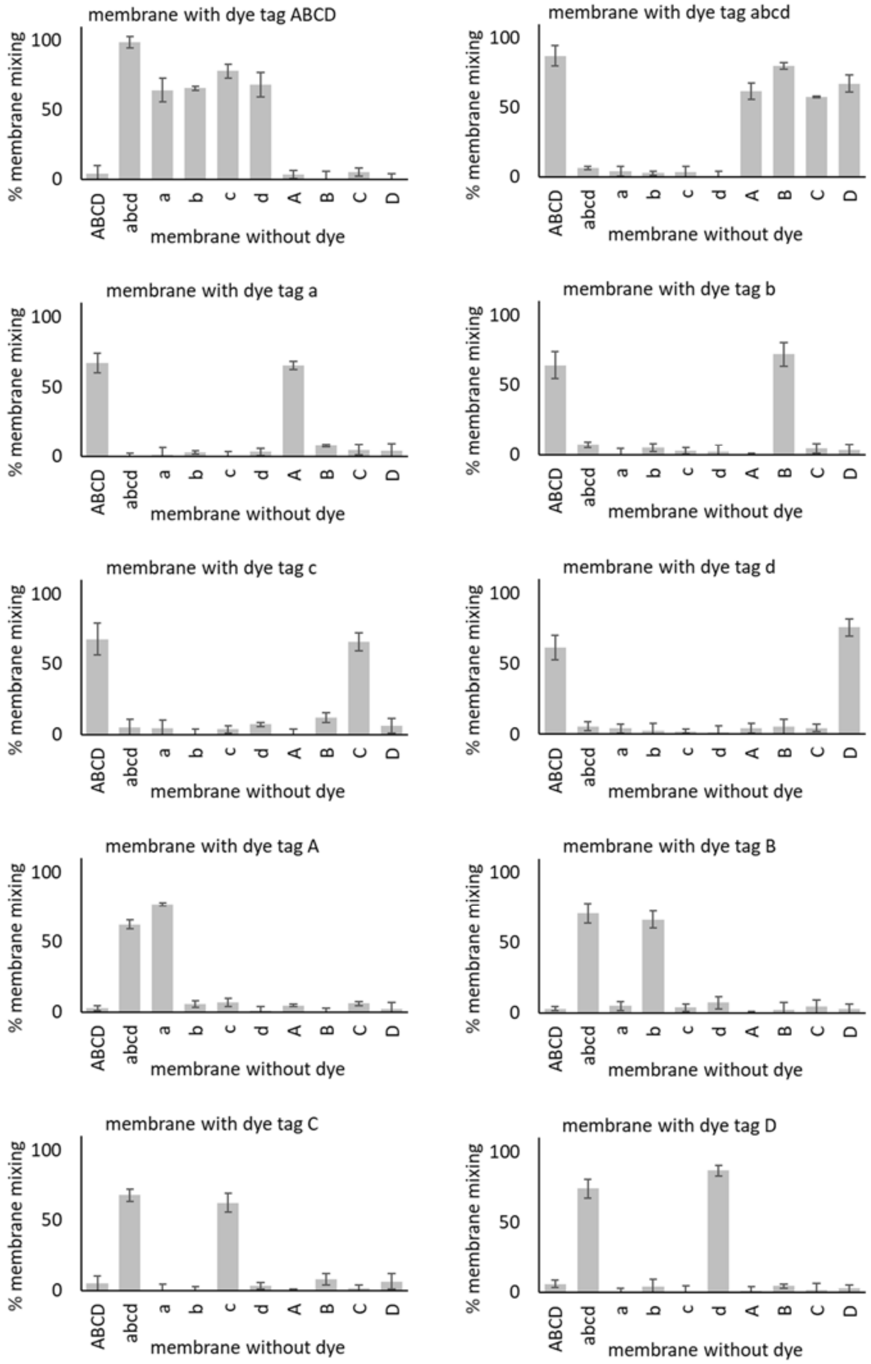


**Figure S28. Individual data points for membrane mixing experiments for liposomes labeled with two different DNA dyes.**

Liposomes with and without FRET dye pairs were prepared with either all (ABCD, abcd) or one of the complementary DNA oligos (a, b, c, d, A, B, C or D). Liposomes were mixed in equimolar amounts, incubated for 1h, and fluorescence was measured. Value represented membrane mixing calibrated from FRET signal (calibration see **Fig. S31**) Error bars indicate +-S.E.M., n=3.

### Figure S29

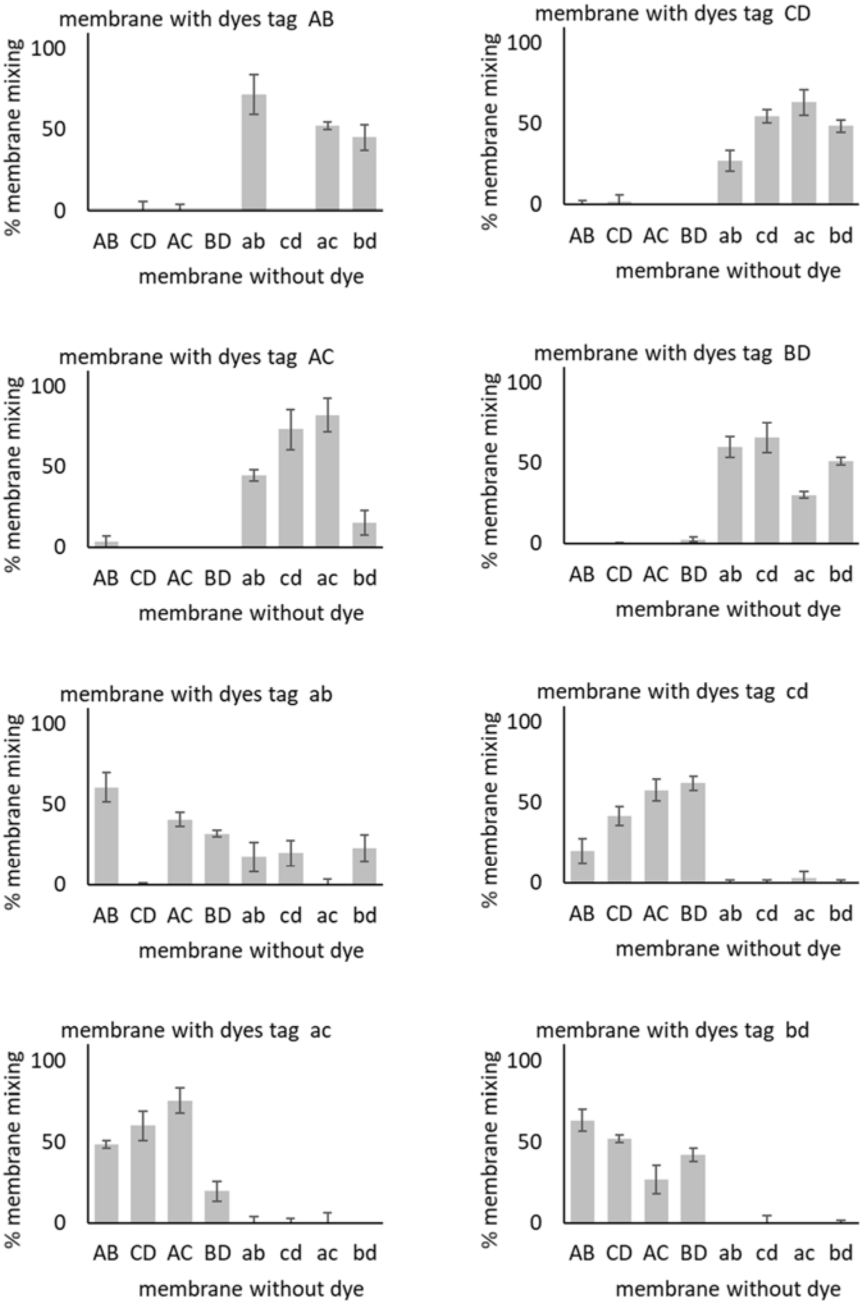


**Figure S29. Individual data points for membrane mixing experiments for liposomes labeled with two different DNA dyes.** Value represented membrane mixing calibrated from FRET signal (calibration see **Fig. S31**) Error bars indicate +-S.E.M., n=3.

### Figure S30

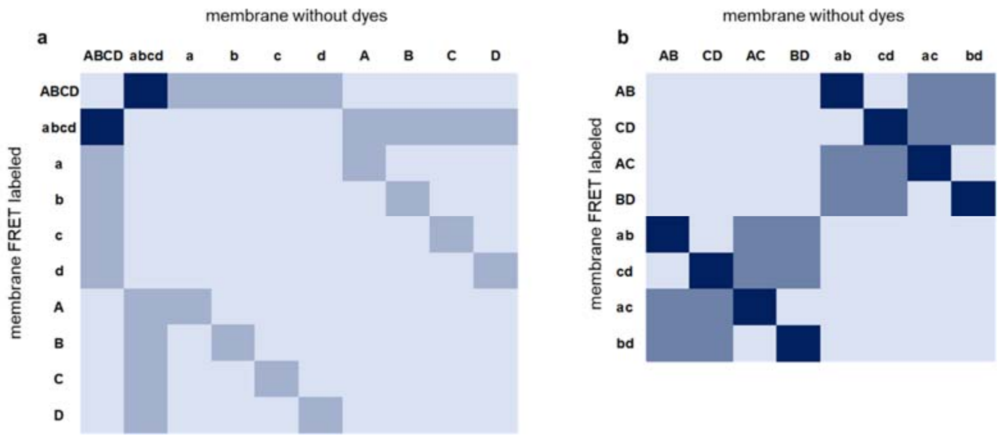


**Figure S30 Theoretically predicted liposome fusion patterns.**
Each experiment assumes mixing of two populations of liposomes, one decorated with complementary pair of membrane-anchored FRET dyes and the other population without any dye. Only if complementary pair (A and a, B and B, C and c, or D and d) is present, the membranes mix. **a**: liposome with FRET dye pairs labeled with all four, or every single one of the possible DNA strands, mixed with liposomes with unlabeled. **b**: same conditions as a, but liposomes labeled with pairs of DNA.
The prediction of mixing efficiency was made by assigning value of 0 if no complementary tag pairs are present on either population, and value of 1 for each pair of complementary tags (A-a, B-b, C-c or D-d), for the maximum value of 4 if ABCD population was mixed with abcd.
The actual experimental data for this system are on **Figure S4**.

### Figure S31

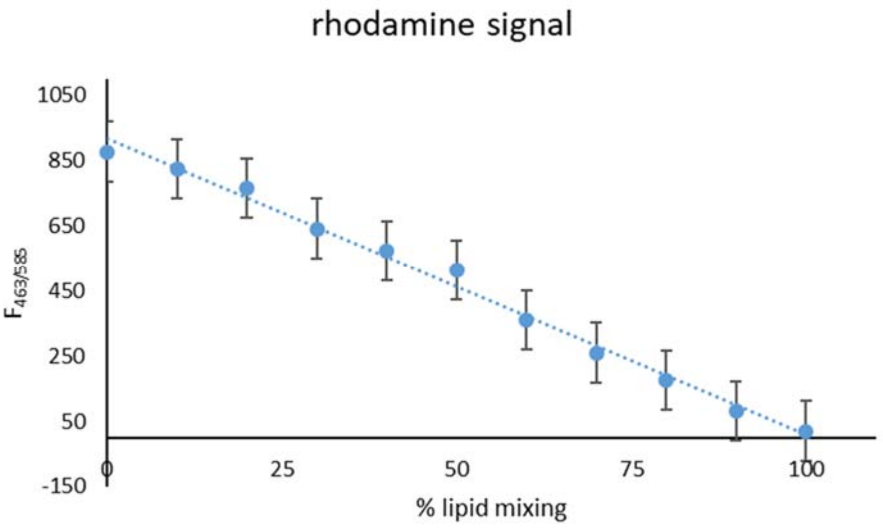


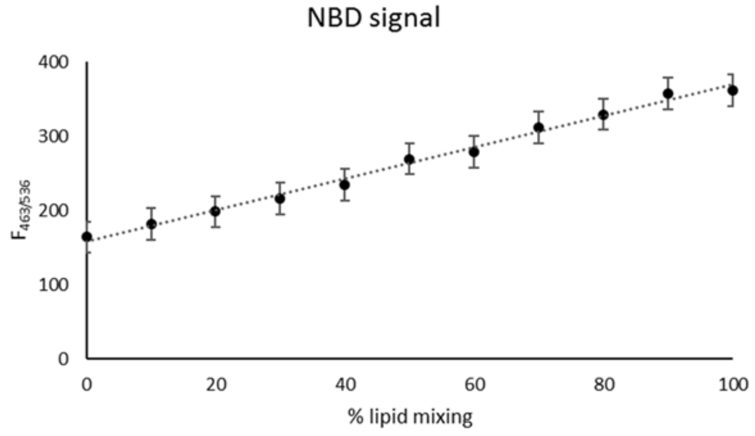


**Figure S31. Calibration of FRET signal for membrane mixing experiments.**
Liposomes were prepared with varying amount of the FRET dye pair, to simulate lipid to dye ratio of corresponding percentage of membrane mixing, according to previously published protocols.[2–4]

### Table S1

| | | |
|---|---|---|
| P28 | Promoter of the tar gene (*E. coli*) specific to σ28 | GenBank: U00096.2, extensive characterization in cell-free Tx/Tl[5] |
| P70 | Lambda phage promoter specific to *E. coli* σ70 (OR2-OR1-Pr) | GenBank: J02459.1, extensive characterization in cell-free Tx/Tl[5] |
| T7 | Promoter of bacteriophage T7 | Genbank: NC_003298 |
| T3 | Promoter of bacteriophage T3 | Genbank: NC_001604 |
| Dre | Dre recombinase | |
| Vika | Vika recombinase | |
| Cre | Cre recombinase | |
| Cre1, Cre2 | split Cre recombinase | |
| T7RNAP1, T&RNAP2 | Split T7 RNA polymerase | |
| Pum | Pumby proteins, based on Pumilio homology domain | Building blocks gift from Ed Boyden, Pumby assembly reference[6] |
| GFP | deGFP, engineered for cell-free bacterial expression | gift from Vincent Noireaux, reference[7] |
| mVenus | mVenus, engineered for cell-free bacterial expression | gift from Vincent Noireaux |
| mApple | mApple, engineered for cell-free bacterial expression | gift from Vincent Noireaux |
| CFP | Cyan fluorescent protein, engineered for cell-free bacterial expression | gift from Vincent Noireaux, reference[7] |
| T7 RNAP | T7 bacteriophage RNA polymerase | gift from Vincent Noireaux, extensive characterization in cell-free Tx/Tl[5] |
| T3 RNAP | T3 bacteriophage RNA polymerase | extensive characterization in cell-free Tx/Tl[5] |
| σ28 | rpoF, E. coli σ28 | GenBank: U00096.2, extensive characterization in cell-free Tx/Tl[5] |
| P28 | binding site for sigma factor 28 | extensive characterization in cell-free Tx/Tl[5] |
| tetR | Tet operon regulatory gene | gift from Ed Boyden, GenBank: BAG71042.1, extensive characterization in cell-free Tx/Tl[5] |
| fLuc | firefly luciferase, utilizing substrate luciferin | gift from Ed Boyden |
| fLucA, fLucB, Scaffold | split firefly luciferase on coiled-coil protein scaffold | gift from Ed Boyden, first published[1] |
| cLuc | *Cypridina* luciferase, utilizing substrate vargulin | GenBank: AB262361.1 |
| rLuc | *Renilla* luciferase, utilizing substrate | gift from Ed Boyden |
| Vio A, Vio B, Vio C, Vio D | Violacein pathway genes | gift from Claudia Schmidt-Dannert, used in cell-free in solution[8] and in lyophilized system[9] |

Table S1 Genes and promoters list.

### Table S2

| | |
|---|---|
| Liposome Fusion Signal Sequence 1 Top Strand | GGT GTC AGT AAG C |
| Liposome Fusion Signal Sequence 1 Bottom Strand | complement to above |
| Liposome fusion signal sequence 2 top strand | CATTCGAGATCCT |
| Liposome fusion signal sequence 2 bottom strand | complement to above |
| Liposome fusion signal sequence 3 top strand | CATAGTCGTCTCA |
| Liposome fusion signal sequence 3 bottom strand | complement to above |
| Liposome fusion signal sequence 4 top strand | CCAGATCCTCATA |
| Liposome fusion signal sequence 4 bottom strand | complement to above |
| Liposome Fusion Sequence 1 Top Strand - chol labeled | GTCTAGCGTCTCACCAG/3CholTEG/ |
| Liposome Fusion Sequence 1 Bottom Strand - chol labeled | CTGGTGAGACGCTAGAC/3CholTEG/ |
| Liposome fusion sequence 2 top strand - chol labeled | ATCCTCATAGTCG/3CholTEG/ |
| Liposome fusion sequence 2 bottom strand - chol labeled | cholesterol labeled complement to above |
| Liposome fusion sequence 3 top strand - chol labeled | TCTCACCAGATCC/3CholTEG/ |
| Liposome fusion sequence 3 bottom strand - chol labeled | cholesterol labeled complement to above |
| Liposome fusion sequence 4 top strand - chol labeled | CTTGGGTTGATTT/3CholTEG/ |
| Liposome fusion sequence 4 bottom strand - chol labeled | cholesterol labeled complement to above |

**Table S2** Complementary ends of strands used in liposome fusion and content mixing experiments. 3CholTEG is 3' Cholesterol-TEG (15 atom triethylene glycol spacer) modification.

### SI Literature


1. Selgrade, D. F., Lohmueller, J. J., Lienert, F. & Silver, P. a. Protein Scaffold-Activated Protein Trans-Splicing in Mammalian Cells. *J. Am. Chem. Soc.* 7713–7719 (2013).
2. Chen, I. a. & Szostak, J. W. A Kinetic Study of the Growth of Fatty Acid Vesicles. *Biophys. J.* **87,** 988–998 (2004).
3. Chen, I. a, Roberts, R. W. & Szostak, J. W. The emergence of competition between model protocells. *Science* **305,** 1474–1476 (2004).
4. Adamala, K., Engelhart, A. E. & Szostak, J. W. Collaboration between primitive cell membranes and soluble catalysts. *Nat. Commun.* **7,** 1–7 (2016).
5. Shin, J. & Noireaux, V. An E. coli cell-free expression toolbox: Application to synthetic gene circuits and artificial cells. *ACS Synth. Biol.* **1,** 29–41 (2012).
6. Adamala, K. P., Martin-Alarcon, D. A. & Boyden, E. S. Programmable RNA-binding protein composed of repeats of a single modular unit. *Proc. Natl. Acad. Sci.* **19,** E2579–E2588 (2016).
7. Shin, J. & Noireaux, V. Efficient cell-free expression with the endogenous E. Coli RNA polymerase and sigma factor 70. *J. Biol. Eng.* **4,** 8 (2010).
8. Garamella, J., Marshall, R., Rustad, M. & Noireaux, V. The all E. coli TX-TL Toolbox 2.0: a platform for cell-free synthetic biology. *ACS Synth. Biol.* **5,** 344–355 (2016).
9. Pardee, K. *et al.* Portable, On-Demand Biomolecular Manufacturing. *Cell* **167,** 248–259.e12 (2016).
